# Reprogramming M2-polarized patient-derived glioblastoma associated microglia/macrophages via CSF1R inhibition

**DOI:** 10.1101/2022.10.20.511747

**Authors:** Valentina Fermi, Rolf Warta, Carmen Rapp, Maximilian Knoll, Gerhard Jungwirth, Christine Jungk, Philip Dao Trong, Andreas von Deimling, Amir Abdollahi, Andreas Unterberg, Christel Herold-Mende

**Affiliations:** Department of Neurosurgical Research, University Hospital Heidelberg, Im Neuenheimer Feld 400, 69120 Heidelberg, Germany; German Cancer Consortium (DKTK), National Center for Tumor Diseases (NCT), Im Neuenheimer Feld 460, 69120 Heidelberg, Germany; Department of Radiation Oncology, University Hospital of Heidelberg, Im Neuenheimer Feld 400, 69120 Heidelberg, Germany; Clinical Cooperation Unit Radiation Oncology, German Cancer Research Center (DKFZ), Im Neuenheimer Feld 280, 69120 Heidelberg, Germany; Heidelberg Institute for Radiation Oncology (HIRO), University Hospital of Heidelberg, Im Neuenheimer Feld 400, 69120 Heidelberg, Germany; Dept. of Neuropathology, University Hospital Heidelberg, Heidelberg, Germany; Clinical Cooperation Unit Neuropathology, German Cancer Consortium (DKTK), German Cancer Research Center, Heidelberg, Germany

## Abstract

Targeting immunosuppressive and protumorigenic glioblastoma-associated macrophages and microglial cells (GAMs) holds great potential to improve patient outcomes. Although CSF1R has emerged as a promising target to reprogram anti-inflammatory M2-like GAMs, relevant treatment data on human, tumor-educated GAMs and innovative patient-derived 3D tumor organoid models to study the influence on adaptive immunity and the effectiveness of treatment in a complex and entirely autologous setting are largely lacking. We performed a comprehensive phenotypical, transcriptional and functional analysis of primary, patient-derived GAMs upon treatment with the CSF1R-targeting drugs PLX3397, BLZ945, and GW2580. The most effective reprogramming of GAMs was observed upon GW2580 treatment, which led to a downregulation of M2-related markers and signaling pathways, while M1-like markers, phagocytosis, and T-cell killing were substantially increased. Moreover, treatment of patient-derived glioblastoma organoids with GW2580 confirmed successful reprogramming together with reduced tumor cell proliferation, indicating that treatment with GW2580 could be an important pillar in the future therapy of GBM.

## Introduction

Glioblastoma (GBM) is one of the most aggressive tumors, and despite intensive standard therapy consisting of surgery and subsequent combined radiochemotherapy, the overall survival of GBM patients is limited to 12-18 months^1^. By initiating a novel or boosting an existing antitumor immune response, the effectiveness of cancer immunotherapy in many otherwise untreatable cancers ^2^ fueled the hope to also improve GBM treatment by targeting cells of the tumor microenvironment (TME). For instance, monoclonal antibodies against programmed cell death protein 1 (PD1) and cytotoxic T-lymphocyte antigen 4 (CTLA-4) have been approved for the treatment of metastatic melanomas^3,4^. However, the treatment of GBM patients with these antibodies in phase III clinical studies failed to improve survival^5,6,^ and to date, no immunotherapeutic approaches have been successful in GBM treatment. In addition to a low tumor mutational burden of GBM^7^, possible reasons for this failure include an immunosuppressive TME in malignant gliomas resulting in a comparably poor infiltration of effector T cells^8^ and various immunosuppressive mechanisms that counteract the effectiveness of antitumor immunity^9^.

In particular, myeloid cells, such as brain-resident microglia and bone marrow-derived macrophages representing up to 30% of the tumor mass, seem to play a key role in the orchestration of an immunosuppressive microenvironment in malignant gliomas^10,11^. Physiologically, microglia and macrophages are involved in tissue homeostasis by phagocytosis of apoptotic cells and by supporting neurogenesis, synaptic refinement, and axonal growth. Moreover, they are an integral component of the first-line defense and thereby contribute substantially to immune surveillance^12^. However, in the presence of tumor cells, glioblastoma-associated microglia/macrophages (GAMs) have been shown to be re-educated toward a pro-tumorigenic, anti-inflammatory M2-like phenotype^13^. As a consequence, GAMs suppress T-cell function by increasing the activity of arginase-1 (ARG1)^14^, reducing the activation of inducible nitric oxide synthase (iNOS)^15^, upregulating the expression of immune checkpoint ligands such as programmed cell death ligand 1 (PD-L1)^16^ and secreting IL-10, which in turn suppresses the activation of cytotoxic T cells^17^.

Several approaches to inhibit the protumorigenic and immunosuppressive functions of myeloid cells have been investigated in mouse models of various cancer types^18^, including malignant gliomas^12^. Among them, the tyrosine kinase receptor colony stimulating factor-1 receptor (CSF1R), which is expressed on mature macrophages^19^, frequently serves as a target because it induces GAMs to promote GBM tumor invasion through a paracrine loop and seems to be involved in the anti-inflammatory activation of macrophages^20^. Additionally, the expression of its ligand CSF1 positively correlates with malignancy in gliomas and is associated with the acquisition of an M2-like phenotype of GAM^21^. Furthermore, in preclinical studies, CSF1R blockade has shown promising results. For instance, treatment of a mouse glioma model with the small molecule inhibitor (SMI) BLZ945 induced tumor regression and improved survival in tumor-bearing mice^22^. Moreover, the SMI PLX3397 improved the effects of irradiation and prolonged the median survival of xenotransplanted glioblastoma-bearing mice compared to irradiation alone^23^.

However, to date, most studies on CSF1R-targeting drugs have been performed in mouse glioma models, which may not entirely represent patient tumors, or on human nontumor-derived macrophages, neglecting the phenotypical and functional changes in the tumor-educated GAM as well as the contribution of microglial cells as part of the entire myeloid compartment. Furthermore, no direct comparison of drug efficacy has been performed to elucidate the best CSF1R-targeting drugs for future treatment or their effects on T-cell immunity. Therefore, we performed a comparative analysis of CSF1R blockade for three of the most promising SMIs (PLX3397, BLZ945, and GW2580) on GAMs freshly isolated from primary human GBM. We studied their ability to shift GAM polarization from an immunosuppressive to a proinflammatory phenotype at the protein, mRNA, and functional levels. We also investigated whether GAM treatment with CSF1R-targeting SMI may prevent the immunosuppressive influence of GAMs on patient-derived T cells. Finally, the effect of CSF1R blockade was validated in an entirely autologous patient-derived tumor organoid model representing the natural tumor environment of GBM.

## Materials and Methods

### Patient samples

Human IDHwt primary GBM (pGBM) specimens (n = 15 for GAM and n = 4 for tumor organoid treatment) were obtained from patients undergoing surgical resection at the Department of Neurosurgery (Heidelberg, Germany). Corresponding blood samples of patients were obtained intraoperatively. The use of patient material was approved by the Institutional Review Board at the Medical Faculty Heidelberg. Informed consent was obtained from all patients included in the study. The clinical and molecular characteristics of the study sample are listed in Supplementary Table 1.

### PBMC isolation and T-cell sorting

Peripheral blood mononuclear cells (PBMCs) were isolated by Ficoll gradient centrifugation (Biochrom, Berlin, Germany) and stored in liquid nitrogen. T cells were isolated from PBMCs using the Dynabeads® Untouched Human T Cells Kit (Invitrogen, Karlsruhe, Germany) following the manufacturer’s protocol.

### Cell culture conditions

To isolate glioblastoma-associated microglia/macrophages (GAM), freshly resected tumor tissues were manually minced with sterile scissors, rinsed through sieves (Becton-Dickinson, Heidelberg, Germany) and further dissociated in HBSS (Biochrom, Berlin, Germany) containing an enzyme mix of DNase I (Sigma‒Aldrich, Taufkirchen, Germany) and Liberase DH Research Grade (Roche, Darmstadt, Germany) using the gentleMACS™ Dissociator (Miltenyi Biotec, Bergisch Gladbach, Germany). CD11b^+^ cells were sorted using CD11b (Microglia) MicroBeads, human and mouse (Miltenyi Biotech, Bergisch Gladbach, Germany) following the manufacturer’s instructions. CD11b^+^ cells were cultivated in DMEM/HAM’s F-12 medium (Invitrogen, Karlsruhe, Germany) containing 20% BIT serum-free supplement (Pelo-Biotech, Plannegg, Germany), 20 ng/ml bFGF, and 20 ng/ml EGF (both Novoprotein, Summit, USA). Glioblastoma-conditioned medium (GCM) was added at a 1:1 ratio.

Tumor-supplying endothelial cells were isolated from single cell suspensions by magnetic bead sorting of CD31^+^ cells using Dynabeads® CD31 Endothelial Cells (Invitrogen, Karlsruhe, Germany) following the manufacturer’s protocol. Cells were cultivated in complete endothelial cell growth medium MV (PromoCell, Heidelberg, Germany) supplemented with 10% human AB serum (Milan Analytica, Rheinfelden, Switzerland).

Tumor cell cultures were generated by cultivation of CD11b-neg and CD31-neg cells in DMEM (Invitrogen, Karlsruhe, Germany) supplemented with 10% fetal calf serum (FCS) (Biochron, Berlin, Germany) and 1% penicillin/streptomycin (Invitrogen, Karlsruhe, Germany). Cell cultures were authenticated by short tandem repeat (STR) DNA profiling analyses (Leibniz Institute DMSZ, Braunschweig, Germany).

### Preparation of glioblastoma-conditioned medium

To prepare glioblastoma-conditioned medium (GCM), tumor cells were seeded in tissue culture flasks. After 24 hours, the medium was changed, and the cells were cultivated for 48 hours in FCS-free DMEM. The supernatant was collected, centrifuged at 4500xg for 10 min at 4 °C and stored at −80 °C. Tumor cells were detached and counted.

### Acetylated low-density lipoprotein (acLDL) uptake test

CD11b^+^ cells were incubated with AlexaFluor® (AF)488-coupled acLDL particles (Invitrogen, Karlsruhe, Germany) diluted 1:500 in culture medium for 4 hours at 37 °C. The supernatant was removed, and the cells were washed twice with PBS to remove free-floating acLDL particles. Internalization of AF488-coupled acLDL particles by CD11b+ cells was monitored using an IX51 fluorescence microscope (Olympus, Hamburg, Germany).

### GAM treatment

The CSF1R-targeting small molecule inhibitors (SMIs) BLZ945, GW2580, and PLX3397 (Selleckchem, Houston, USA) were reconstituted in DMSO at stock concentrations of 1 or 10 mM. For flow cytometry and transcriptional analyses, CD11b^+^ cells were seeded in T25 culture flasks at a density of 100,000 cells/ml in 3 ml culture medium. Treatment with SMIs started on the next day. SMIs were diluted in culture medium to a final concentration of 1 μM. Control cells were incubated with DMSO (0.1%). Treatment was performed in total for 96 hours, while after 48 hours, the medium was changed to DMEM/HAM F-12 medium containing the respective SMI, BIT supplement, bFGF, and EGF without GCM.

### Formation, cultivation, and treatment of tumor organoids

To generate tumor organoids, freshly prepared tumor-derived single-cell suspensions were incubated overnight with NanoShuttle™-PL (Greiner Bio-One, Kremsmünster, Austria) at a concentration of 1 μl NanoShuttle/10,000 cells. Thereafter, cells were seeded in a CELLSTAR® Cell-Repellent 96-well plate (Greiner Bio-One, Kremsmünster, Austria) at a density of 50,000 cells per well in 150 μl culture medium. The plate was placed on top of a spheroid drive (Greiner Bio-One, Kremsmünster, Austria) for 15 min at 37 °C to let cells aggregate at the bottom of each well. Tumor organoids were cultivated in DMEM/HAM F-12 medium containing BIT supplement, bFGF, and EGF for 5 days. The medium was partially replaced on day 3. Cell viability in tumor organoids was monitored with the LIVE/DEAD™ Cell Imaging Kit (Invitrogen, Karlsruhe, Germany) according to the manufacturer’s instructions. After 5 days, tumor organoids were treated with SMI at a concentration of 10 μM, while incubation with DMSO (0.1%) served as a control. After 96 hours, tumor organoids were shock-frozen in precooled 2-methylbutan (Carl Roth, Karlsruhe, Germany) using a magnetic pen (Greiner Bio-One, Kremsmünster, Austria) to transfer them into cryovials.

### Flow cytometry

CD11b^+^ cells were detached by incubation with 0.25% Trypsin/EDTA (Invitrogen, Karlsruhe, Germany) for 2 min at 37 °C and stained using the LIVE/DEAD™ Fixable Violet Dead Cell Stain Kit (Thermo Fisher Scientific, Waltham, USA) according to the manufacturer’s protocol. Next, the cells were incubated for 20 min at RT with an anti-human CD163-PE antibody and an anti-human HLA-DR-PE-Cy7 antibody according to the manufacturer’s protocol. Antibodies and the LIVE/DEAD Fixable Stain Kit were diluted in PBS containing 0.5% human serum and 2 mM EDTA. For labeling of intracellular proteins such as CD68, the FoxP3/Transcription Factor Staining Buffer Set (Thermo Fisher Scientific, Waltham, USA) was used for fixation and permeabilization according to the manufacturer’s instructions. Then, the cells were stained with an anti-human CD68-FITC antibody diluted in the permeabilization buffer of the FoxP3/Transcription Factor Staining Buffer Set for 30 min at 4 °C. Detailed information about the antibodies can be found in Supplementary Table 2. Cells were analyzed by flow cytometry using an LSRII flow cytometer (Becton-Dickinson, Heidelberg, Germany), FACSDiva software (Becton-Dickinson, Heidelberg, Germany), and FlowJo Software v10.7.1 (TreeStar, San Carlo, USA).

### Multicolor immunofluorescence staining

Slides with two consecutive sections of tumor organoids were stained with antibodies against GFAP, TNC, Ki-67, CD68, CD163, and CD3. All primary antibodies were diluted with Antibody Diluent (Dako, Hamburg, Germany) and incubated for 1 hour at RT. To counterstain nuclei, DAPI (Thermo Fisher Scientific, Waltham, USA) was diluted 1:1000 in DPBS. Appropriate fluorochrome-conjugated secondary antibodies were diluted in DAPI solution and incubated for 1 hour in the dark. When using two primary antibodies from the same species, one of them was directly coupled with a fluorophore using the Zenon Labeling Kit (Thermo Fisher Scientific, Waltham, USA) according to the manufacturer’s instructions and then incubated for 20 min (for anti-CD68 antibody) or 1 hour (for anti-Ki-67 antibody) in the dark. Three washing steps with PBS containing 0.05% Tween (Sigma‒Aldrich, Taufkirchen, Germany) were performed after the application of primary and secondary antibodies. Finally, slides were mounted with Elvanol (Roth, Karlsruhe, Germany), and a fluorescence microscope IX51 equipped with an F-View II camera (Olympus, Hamburg, Germany) was used to image fluorescent signals. Detailed information about the primary and secondary antibodies can be found in Supplementary Tables 3 and 4, respectively. Images were analyzed and quantified using CellSense Dimension (Olympus, Hamburg, Germany) and StrataQuest Software (TissueGnostic, Vienna, Austria).

### Hematoxylin-eosin staining

Cryosections of tumor organoids were incubated with hematoxylin solution (Carl Roth, Karlsruhe, Germany) for 1 min and then washed with tap water. Next, incubation with the eosin G solution (Carl Roth, Karlsruhe, Germany) was performed for 5 seconds, followed by a dehydration procedure with an increasing percentage of ethanol (70%, 90%, and 100%). Then, the slides were immersed twice in 100% xylene (Carl Roth, Karlsruhe, Germany) and dried overnight before adding CytosealTM 60 (Thermo Fisher Scientific, Waltham, USA) and a coverslip.

### RNA isolation and RNA sequencing

RNA was isolated using the fully automated sample preparator QIAcube in combination with the QIAgen RNeasy® Micro Kit (Qiagen, Hilden, Germany). The RNA concentration was measured using a Nanodrop 2000 spectrophotometer (Thermo Fisher Scientific, Waltham, USA). RNA quality and integrity were tested with a Bioanalyzer 2100 (Agilent Technologies, Santa Clara, USA). Transcriptome analyses were performed at the Genomic and Proteomics Core Facility of the German Cancer Research Center (Heidelberg, Germany). RNA libraries (low input, NEBNext) were sequenced on an Illumina HiSeq 4000 system. Reads were mapped to the human reference genome GRCh38.p7 and quantified with STAR v. 2.6.0c^24^. Analysis and identification of differentially expressed genes was performed by DESeq2 (3.15) in R (4.0).

### Phagocytosis assay

CD11b^+^ cells were stained using the Cell Proliferation Dye eFluor™ 670 (Thermo Fisher Scientific, Waltham, USA) according to the manufacturer’s instructions and seeded in a flat-bottom 96-well plate at a density of 100,000 cells per well in 100 μl culture medium. After overnight incubation, the cells were treated with SMIs as described. After treatment, the culture medium was replaced by 100 μl of pHrodo™ E. coli BioParticles® Conjugates (Thermo Fisher Scientific, Waltham, USA) suspension, prepared following the manufacturer’s instructions. After 4 hours of incubation at 37 °C, the cells were detached as described, transferred to a 96-well plate with U-bottom, and analyzed by flow cytometry using an LSRII flow cytometer, FACSDiva software and FlowJo Software v10.7.1.

### Griess diazotization reaction-based assay

After treatment of CD11b+ cells, the supernatant was collected and centrifuged at 4500xg for 10 min at 4 °C. GAM-conditioned medium was stored at −80 °C until further use. After thawing, the nitrite concentration was measured in GAM-conditioned media by a Griess diazotization reaction-based assay (Thermo Fisher Scientific, Waltham, USA) following the manufacturer’s protocol at an absorbance of 548 nm using an Infinite® 200 photometer (Tecan, Männerdorf, Switzerland).

### Bead-based immune assay

To measure the concentrations of cytokines, chemokines, and growth factors in GAM-conditioned media, Luminex analyses were performed using the Bio-Plex ProTM Human Cytokine 27-Plex Assay as well as the Bio-Plex Pro™ TGF-β Assay (both Bio-Rad Laboratories, Hercules, USA) according to the manufacturer’s protocols on the Bio-Plex® Reader 200 by using the Bio-Plex Manager™ software.

### Transmigration assay

To test the transmigration of T cells through a dense barrier of tumor-derived endothelial cells, 24-well plates and ThinCert™ Cell Culture Inserts (Greiner Bio-One, Kremsmünster, Austria) with a pore diameter of 3 μm were used to divide each well into upper and lower chambers. Cell culture inserts were precoated with 100 μl gelatin solution (Sigma‒Aldrich, Taufkirchen, Germany) for 30 min at 37 °C before seeding tumor-derived endothelial cells (30,000 cells/insert; 100 μl endothelial cell growth medium MV-2, PromoCell Heidelberg, Germany). Then, 500 μl of culture medium was added to each lower chamber. Endothelial cells grew as monolayers on the inserts for four days. The tightness of the monolayer was tested by performing an Evans blue dye exclusion test. Endothelial cells were incubated with GAM-conditioned media diluted 1:10 in culture medium for 48 hours. T cells were isolated from PBMCs as described and cultivated overnight in X-Vivo 20 (Lonza, Basel, Switzerland) supplemented with 10% human serum, IL-2 (100 IU/ml) (Novartis, Basel, Switzerland), and IL-4 (60 IU/ml) (Miltenyi Biotech, Bergisch Gladbach, Germany). T cells were added to the upper chamber (30,000 cells/chamber in 100 μl MV-2 medium). After 24 hours, the supernatant in the lower chamber was collected, and the number of transmigrated T cells was measured using an LSRII flow cytometer in a 96-well plate format by using FACSDiva software and FlowJo Software v10.7.1.

### Evans blue dye exclusion test

To perform the Evans blue dye exclusion test, stock solution (Sigma‒Aldrich, Taufkirchen, Germany) was diluted 1:10 in Endothelial Cell Basal Medium MV (PromoCell, Heidelberg, Germany). Ten microliters of the diluted dye was added to each upper chamber and incubated for 20 min at RT. The lower chambers were optically investigated for the presence of dye traces. A monolayer of endothelial cells was considered tight when no traces of Evans blue dye were detectable in the lower chamber.

### Immune cell killing assay

Immune cell killing assays were performed to investigate the effects of GAM treatment on the ability of T cells to kill autologous tumor cells. Tumor cells were stained with CellTrace™ CFSE (Thermo Fisher Scientific, Waltham, USA) according to the manufacturer’s instructions. Next, they were seeded in a flat-bottom 96-well plate at a density of 5,000 cells/well in 100 μl culture medium and cultivated overnight at 37 °C. T cells were isolated from PBMCs as described and added to tumor cells at a 1:10 target-to-effector ratio. Tumor and T cells were incubated with GAM-conditioned medium diluted 1:20 in X-Vivo 20 supplemented with 10% human serum for 48 hours. Then, the cells were collected, stained with propidium iodide (Thermo Fisher Scientific, Waltham, USA) following the manufacturer’s protocol, and counted using an LSRII flow cytometer in a 96-well plate, FACSDiva software, and FlowJo Software v10.7.1.

### Live-cell imaging of tumor cell-T-cell cocultures

Tumor cells were seeded in a flat-bottom 96-well plate at a density of 5,000 cells/well in 100 μl culture medium and cultivated overnight at 37 °C. Then, they were stained with the red-fluorescent dye MytoTracker™ (Thermo Fisher Scientific, Waltham, USA) following the manufacturer’s instructions. T cells were isolated from PBMCs as described and added to tumor cells at a 1:10 target-to-effector ratio. The green fluorescent IncuCyte™ Caspase-3/7 Apoptosis Reagent (Essen BioScience, Ann Arbor, USA) was added at a concentration of 20 μM. Tumor and T cells were incubated with GAM-conditioned media diluted 1:20 in X-Vivo 20 supplemented with 10% human serum for 48 hours. The IncuCyte™ Zoom Instrument (Essen BioScience, Ann Arbor, USA) was used for live cell imaging. Images of whole wells were taken with a 4X objective and by using a combination of three channels: red, green, and phase contrast. The plate was scanned every two hours, and 4 images per well were taken at each scan session.

### Analysis of TCGA data

TCGA (The Cancer Genome Atlas) data were obtained using the web application GlioVis (http://www.gliovis.bioinfo.cnio.es). Microarray data of n = 528 pGBM and 10 normal brain tissues were used for expression analysis. For survival analysis, n = 357 IDHwt pGBM and corresponding clinical data were used.

### Statistical analysis

Data were analyzed using GraphPad Prism 9.02 software and the R software environment (4.0). For all analyses, a two-sided Student’s t test was used. A p value < 0.05; **p < 0.01; ***p < 0.001 is considered to be significant. Error bars show the standard error of the mean (SEM) of technical triplicates. A log-rank test was performed to compare the survival data of the two groups (high vs. low CSF1R expression, divided by the median), and Kaplan‒Meier curves were used to visualize survival data.

## Abbreviations

GAM: Glioblastoma-associated microglia/macrophages
IDH: isocitrate dehydrogenase (NADP(+))
MGMT: O-6-methylguanine-DNA methyltransferase
CSF1R: colony stimulating factor-1 receptor
GBM: glioblastoma
ARG1: arginase-1
iNOS: inducible nitric oxide synthase
SMI: small molecule inhibitor

## Data availability

RNA sequencing data have been deposited in NCBI’s Gene Expression Omnibus (GEO). Data for GAM upon treatment with CSF1R inhibitors was deposited under accession code “soon available”, and tumor organoids upon treatment with CSF1R inhibitors was deposited under accession code “soon available”. Source data are provided with this paper.

## Results

### CSF1R as a potential target in GBM

There is increasing evidence in several tumor types, which indicates that CSF1R is a promising target to change the activation status of anti-inflammatory M2-like GAMs^12,25^. When analyzing the expression of CSF1R in a dataset of 528 pGBM samples from the TCGA, we observed significantly increased expression of the CSF1R gene in GBM samples compared to nontumor samples (p = 0.0189) (Fig. 1A). Moreover, higher CSF1R expression in IDHwt pGBM was associated with poorer patient survival (p = 0.043, Fig. 1B) and with the expression of CD163, a scavenger receptor characteristic for M2-like GAM (r = 0.6714, p < 0.0001, Fig. 1C)^26^, corroborating an important role of CSF1R in the polarization of GAMs toward a protumorigenic phenotype and exploiting it as a promising target to modulate the activation of GAMs.

**Figure 1:**
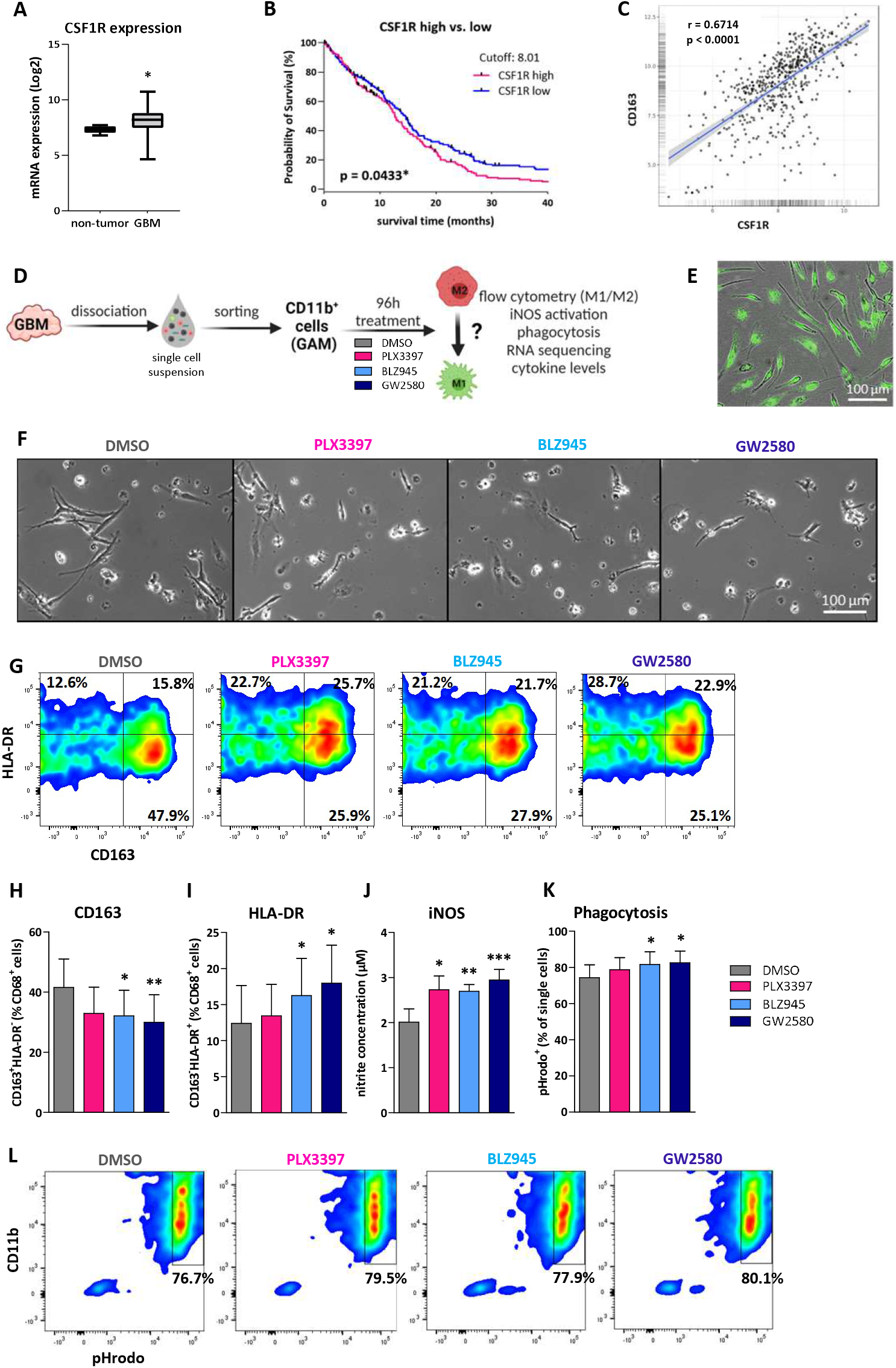
CSF1R targeting of patient-derived GAM results in marked phenotypic and functional M1-like changes. **A** Expression level of CSF1R in pGBM determined in the TCGA microarray data (n=528) compared to nontumor samples (n=10). **B** Survival analysis of IDHwt pGBM patients (n=357) with high (pink) and low (blue) CSF1R expression levels based on TCGA microarray data. **C** Spearman’s rank correlation of CSF1R and CD163 expression in pGBM samples (n=528). **D** Workflow of GAM isolation, treatment and characterization. **E** Microscopic image of a pure GAM culture as determined by acLDL uptake. **F** Microscopic images of GAM morphology upon SMI treatment. **G** Flow cytometry plots illustrating GAM expression of CD163 and HLA-DR in control (DMSO) and SMI-treated CD11b+ cells. **H** and **I** show the percentages of CD163+HLA-DR- and CD163-HLA-DR+ GAM, respectively, as determined by flow cytometry (n = 6). **J** Nitrite concentration measured in conditioned media of SMI-treated and untreated GAM to assess iNOS activation (n = 10). **K** Percentage of phagocytizing GAM (pHrodo+) upon CSF1R blockade, as assessed by flow cytometry (n = 6). **L** Flow cytometry plots illustrating the percentage of pHrodo+ GAM after phagocytosis of pHrodo™ *E. coli* BioParticles® conjugates. Figures show the standard error of the mean. * p < 0.05; ** p < 0.01; *** p < 0.001; *E. coli* = *Escherichia coli*.

### Activation changes in pGBM patient-derived GAMs upon CSF1R blockade

We next investigated whether blocking CSF1R using PLX3397, BLZ945 or GW2580 can reprogram M2-like GAMs by reducing their anti-inflammatory and protumorigenic activation. To this end, patient-derived GAMs were isolated by magnetic sorting of CD11b^+^ cells from single-cell suspensions prepared from freshly resected IDHwt pGBM specimens (Fig. 1D). The purity of GAM cultures was controlled by the ability to internalize fluorescent-coupled acLDL particles (Fig. 1E).

Next, the GAMs were treated with SMIs at a concentration of 1 μM compared to the DMSO control. Upon CSF1R inhibition, cell morphology changed to less elongated and more rounded, amoeboid cells (Fig. 1F). Moreover, as a general trend for all SMIs used, the expression of the M2 marker CD163 decreased, while the expression of the M1 marker HLA-DR increased (Fig. 1G). However, only treatment with BLZ945 and GW2580 resulted in a significantly reduced percentage of CD163^+^HLA-DR^−^ GAM (p < 0.05 and 0.01, respectively) (Fig. 1H) and an increased percentage of CD163^−^HLA-DR^+^ GAM (p < 0.05) (Fig. 1I).

Furthermore, we assessed the nitrite concentration in GAM-conditioned media upon treatment with the CSF1R-targeting SMI as an indirect measurement of iNOS activation. Compared to the DMSO control, the nitrite concentrations in GAM-conditioned media of PLX3397-(p < 0.05), BLZ945-(p < 0.01), and GW2580-treated GAM (p < 0.001) were significantly increased (Fig. 1J). To further determine the phagocytic ability of GAM upon CSF1R blockade, we incubated SMI-treated GAM with pHrodo™ *E. coli* BioParticles® conjugates and measured the percentage of phagocytizing GAM (CD11b^+^pHrodo^+^ cells) by flow cytometry. Again, only for BLZ945 and GW2580 did we observe a significantly increased ability to phagocytize fluorescent *E. coli* particles compared to untreated cells (Figure 1K-L).

Altogether, targeting CSF1R, especially with GW2580 and BLZ945, was able to shift GAM activation toward a proinflammatory M1-like phenotype and to increase their phagocytic activity.

### Treatment-related transcriptional changes in patient-derived GAM

We next assessed changes in the transcriptional profile upon treatment of GAM with CSF1R inhibitors. A total of 401 genes were differentially expressed between SMI-treated and untreated GAMs (adj. p value < 0.05, Log2(FC) > 0.5) (Fig. 2A-B, Tables 1 & 2, Suppl. Table 4 & 6). Most of them were shared by at least two inhibitors, with 150 genes shared by all three inhibitors. Among them, we observed downregulated expression of the M2 marker CD163, matrix metalloproteinase 9 (MMP9), and S100 calcium binding protein A9 (S100A9), which have both been shown to be poor prognosticators in glioma^27,28^. In contrast, upregulated expression of CD74 is a good prognosticator in GBM^29^, member 14 of the tumor necrosis factor superfamily (TNFSF14) sensitizes tumor cells to IFNγ-induced apoptosis^30^, and M1-related C-C motif chemokine receptor 7 (CCR7)^31^ was shared by all three SMIs (Fig. 2A, Table 2, Suppl. Table 6). BLZ945 and GW2580 additionally reduced the expression of another M2 marker, mannose receptor C-type 1 (MRC1), and the cytokine IL-6, which is regarded as tumor-supportive^32^ (Fig. 2C, Suppl. Fig. S1A, Table 1, Suppl. Table 5). Among the three inhibitors, the most remarkable transcriptional changes were observed for GW2580, which induced the downregulation of additional genes involved in cancer survival, progression, and invasion, such as C-C motif chemokine ligand 13 (CCL13)^33^, CD38^34^, C-X-C motif chemokine ligand 1 (CXCL1)^35^, integrin subunit alpha 4 (ITGA4) and secreted phosphoprotein 1 (SPP1)^36^. Furthermore, gene set enrichment analyses revealed that treatment of GAMs with GW2580 resulted in lower levels of transcripts related to IL10 signaling and the IL6-STAT3 signaling pathway compared to untreated cells (Fig. 2D, Table 2, Suppl. Table 6). Protein quantification confirmed a significantly reduced secretion of the tumor-supportive cytokine IL-6 in GAMs upon treatment in GAM-conditioned medium (p < 0.05) (Fig. 2E). In addition, “antigen processing and presentation of peptide antigen” appeared among the 20 top differentially expressed GO terms in GW2580- and BLZ945-treated GAM compared to untreated controls (Suppl. Fig. S2).

**Figure 2:**
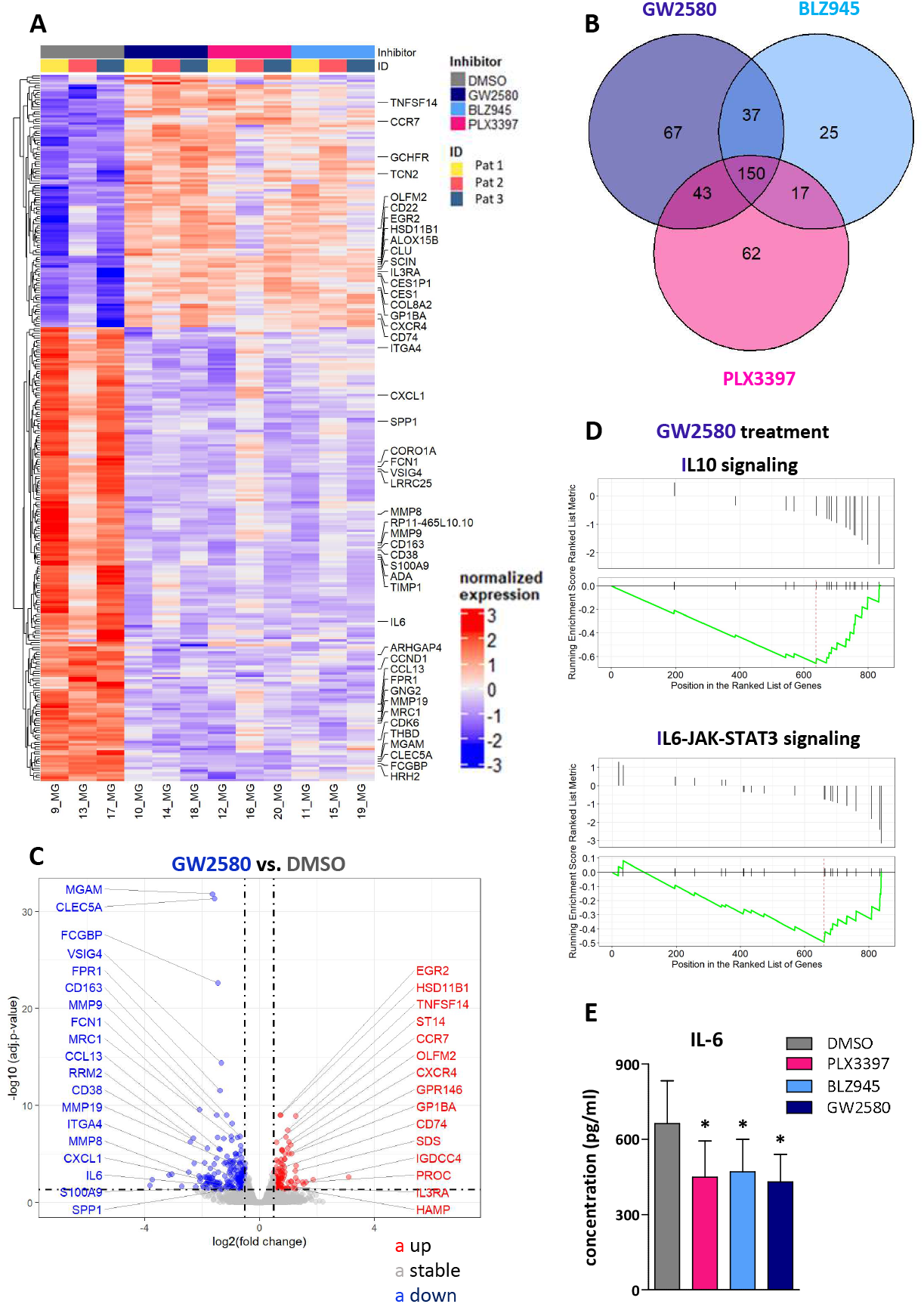
Transcriptional changes in SMI-treated GAMs corroborate M1-like reprogramming. **A** Heatmap of differentially expressed genes upon treatment with GW2580 compared to untreated, BLZ945-treated and PLX3397-treated GAMs (n = 3). **B** Venn diagram showing the overlap of differentially expressed genes. **C** Volcano plot of differentially expressed genes in GW2580-treated GAMs compared to untreated GAMs. Blue = downregulated, red = upregulated genes. Level of significance: adjusted p value < 0.05, Log2(FC) > 0.5. **D** Gene set enrichment analyses showing significantly reduced levels of transcripts related to IL10 signaling and the IL6-STAT3 pathway upon GAM treatment with GW2580. **E** IL-6 protein levels in conditioned media of SMI-treated and untreated GAM (n = 8). Whiskers depict SEM. * p < 0.05; Log = logarithm; FC = fold change; adj. = adjusted; SEM = standard error of the mean.

**Table 1:** Top 30 downregulated genes upon GW2580-based CSF1R blockade in pGBM patient-derived GAM. Abbreviations: CSF1R = colony stimulating factor-1 receptor; pGBM = primary glioblastoma; GAM = glioblastoma-associated microglia/macrophages; Log = Logarithm; FC = fold change; Adj. = adjusted

| Gene | GW2580 |  | BLZ945 |  | PLX3397 |  |
| --- | --- | --- | --- | --- | --- | --- |
|  | Adj. p-value | Log2(FC) | Adj. p-value | Log2(FC) | Adj. p-value | Log2(FC) |
| MGAM | 1.54E-32 | -1.65 | 1.04E-25 | -1.46 | 2.35E-40 | -1.82 |
| CLEC5A | 4.79E-32 | -1.57 | 5.35E-28 | -1.48 | 2.35E-40 | -1.75 |
| FCGBP | 2.51E-23 | -1.45 | 1.68E-21 | -1.40 | 3.86E-26 | -1.53 |
| VSIG4 | 4.09E-15 | -1.34 | 1.32E-14 | -1.32 | 1.82E-15 | -1.35 |
| FPR1 | 3.00E-12 | -1.38 | 2.25E-11 | -1.33 | 1.28E-11 | -1.33 |
| FCN1 | 2.77E-10 | -2.10 | 6.00E-08 | -1.82 | 8.57E-10 | -2.00 |
| CD163 | 1.03E-09 | -1.15 | 6.00E-08 | -1.05 | 2.67E-13 | -1.32 |
| MT1P3 | 1.03E-09 | -1.49 | 6.96E-10 | -1.50 | 8.69E-06 | -1.15 |
| IL1B | 7.72E-09 | -0.96 | 2.56E-09 | -0.99 | 6.73E-05 | -0.72 |
| MMP9 | 1.67E-07 | -0.67 | 1.49E-07 | -0.67 | 3.43E-11 | -0.80 |
| MRC1 | 1.67E-07 | -0.99 | 5.98E-06 | -0.88 | 2.75E-04 | -0.74 |
| GNG2 | 1.94E-07 | -1.47 | 2.03E-06 | -1.35 | 2.79E-05 | -1.22 |
| CXCL3 | 2.04E-07 | -0.76 | 2.15E-07 | -0.76 | 1.23E-04 | -0.61 |
| RRM2 | 2.42E-07 | -2.31 | 6.44E-07 | -2.24 | 1.77E-09 | -2.59 |
| MT1X | 2.70E-07 | -1.25 | 4.73E-08 | -1.31 | 2.58E-05 | -1.06 |
| MT2A | 4.03E-07 | -0.97 | 2.05E-08 | -1.06 | 7.75E-05 | -0.80 |
| IL1R2 | 6.09E-07 | -2.42 | 9.40E-07 | -2.38 | 2.27E-09 | -2.79 |
| RP11-465L10.10 | 1.39E-06 | -0.67 | 1.40E-06 | -0.67 | 4.59E-10 | -0.81 |
| CD38 | 2.48E-06 | -1.82 | 7.37E-06 | -1.74 | 2.65E-07 | -1.93 |
| IRG1 | 2.77E-06 | -1.41 | 3.63E-03 | -1.00 | 2.38E-04 | -1.16 |
| CCL13 | 3.48E-06 | -1.38 | 1.27E-03 | -1.06 | 8.11E-04 | -1.07 |
| LRRC25 | 4.19E-06 | -0.64 | 4.53E-06 | -0.64 | 8.16E-08 | -0.71 |
| CDK6 | 9.05E-06 | -0.84 | 3.59E-05 | -0.80 | 1.40E-04 | -0.75 |
| STAB1 | 9.79E-06 | -0.82 | 5.03E-03 | -0.61 | 7.48E-10 | -1.06 |
| CCND1 | 1.06E-05 | -0.70 | 2.89E-08 | -0.85 | 2.36E-04 | -0.62 |
| CORO1A | 1.64E-05 | -0.81 | 5.98E-06 | -0.84 | 4.61E-07 | -0.90 |
| FLT1 | 1.64E-05 | -1.12 | 6.21E-06 | -1.17 | 3.16E-04 | -0.98 |
| TMEM106A | 2.63E-05 | -0.62 | 8.22E-04 | -0.53 | 9.05E-05 | -0.59 |
| FPR2 | 2.82E-05 | -1.49 | 1.05E-03 | -1.25 | 1.53E-04 | -1.37 |
| WFDC21P | 4.94E-05 | -1.02 | 6.94E-04 | -0.91 | 1.28E-02 | -0.72 |

**Table 2:** Top 30 upregulated genes upon GW2580-based CSF1R blockade in pGBM patient-derived GAM. Abbreviations: CSF1R = colony stimulating factor-1 receptor; pGBM = primary glioblastoma; GAM = glioblastoma-associated microglia/macrophages; Log = Logarithm; FC = fold change; Adj. = adjusted

| Gene | GW2580 |  | BLZ945 |  | PLX3397 |  |
| --- | --- | --- | --- | --- | --- | --- |
|  | Adj. p-value | Log2(FC) | Adj. p-value | Log2(FC) | Adj. p-value | Log2(FC) |
| HSD11B1 | 1.03E-09 | 0.73 | 4.84E-12 | 0.81 | 2.29E-15 | 0.89 |
| EGR2 | 1.03E-09 | 0.72 | 6.26E-11 | 0.76 | 4.97E-15 | 0.87 |
| TNFSF14 | 1.19E-09 | 1.25 | 1.08E-04 | 0.90 | 2.65E-10 | 1.28 |
| ST14 | 3.52E-08 | 0.97 | 6.00E-04 | 0.69 | 1.33E-07 | 0.93 |
| COL8A2 | 1.74E-07 | 0.93 | 2.06E-06 | 0.86 | 1.11E-06 | 0.87 |
| OLFM2 | 5.72E-07 | 1.03 | 1.02E-05 | 0.93 | 6.05E-04 | 0.78 |
| CCR7 | 6.39E-07 | 0.58 | 6.13E-09 | 0.66 | 1.93E-17 | 0.89 |
| CXCR4 | 1.18E-06 | 1.07 | 5.33E-04 | 0.85 | 9.39E-04 | 0.81 |
| ALOX15B | 2.55E-06 | 0.98 | 4.84E-08 | 1.09 | 1.55E-08 | 1.11 |
| CES1 | 3.73E-06 | 0.77 | 1.78E-05 | 0.73 | 7.86E-06 | 0.75 |
| CES1P1 | 3.92E-06 | 0.79 | 4.67E-05 | 0.73 | 2.58E-05 | 0.74 |
| GCHFR | 4.87E-06 | 0.79 | 1.33E-03 | 0.62 | 1.27E-03 | 0.62 |
| GP1BA | 9.79E-06 | 1.11 | 9.73E-06 | 1.11 | 6.53E-07 | 1.20 |
| GPR146 | 9.79E-06 | 1.02 | 5.06E-03 | 0.76 | 1.29E-03 | 0.82 |
| CEBPA | 9.79E-06 | 0.90 | 1.81E-03 | 0.71 | 3.16E-04 | 0.77 |
| NUPR1 | 3.43E-05 | 0.86 | 8.17E-03 | 0.64 | 6.28E-03 | 0.64 |
| CLU | 3.53E-05 | 0.88 | 7.84E-05 | 0.86 | 7.20E-08 | 1.06 |
| TCN2 | 3.70E-05 | 0.75 | 7.75E-04 | 0.66 | 9.35E-03 | 0.54 |
| TNS1 | 4.02E-05 | 0.59 | 2.92E-03 | 0.48 | 7.13E-03 | 0.44 |
| LSP1 | 7.04E-05 | 0.64 | 7.50E-02 | 0.40 | 4.72E-03 | 0.51 |
| SERPINE1 | 9.14E-05 | 0.75 | 1.67E-01 | 0.42 | 9.70E-03 | 0.57 |
| APOC1P1 | 1.26E-04 | 0.80 | 3.66E-02 | 0.55 | 6.93E-02 | 0.50 |
| DEGS2 | 1.75E-04 | 1.27 | 3.37E-03 | 1.09 | 1.82E-03 | 1.11 |
| KLHL6 | 2.40E-04 | 0.57 | 2.66E-03 | 0.50 | 1.41E-04 | 0.58 |
| SNAI3 | 2.51E-04 | 0.87 | 1.39E-02 | 0.68 | 1.82E-03 | 0.77 |
| APOC1 | 2.68E-04 | 0.77 | 5.17E-02 | 0.53 | 5.80E-02 | 0.51 |
| CD74 | 3.35E-04 | 0.68 | 2.16E-04 | 0.70 | 5.34E-03 | 0.57 |
| SIGLEC8 | 5.32E-04 | 0.79 | 1.83E-03 | 0.74 | 1.03E-01 | 0.50 |
| SDS | 5.74E-04 | 1.24 | 6.85E-03 | 1.08 | 1.42E-01 | 0.75 |
| DHRS3 | 6.06E-04 | 0.70 | 1.00E-02 | 0.59 | 4.72E-02 | 0.49 |

Transcriptional changes substantiated a shift in GAM polarization from a tumor-supportive to an immunostimulatory phenotype under the influence of CSF1R-targeting SMI, particularly for GW2580.

### Impact of GAM treatment on T-cell transmigration through an autologous endothelial cell barrier

We next assessed the additional impact of treatment-related changes in the GAM phenotype and activation on T-cell properties. Due to the frequently disrupted ability of tumor-reactive T cells to pass the blood-tumor barrier^8^, we first studied the transmigration of patient-derived T cells through a barrier of autologous endothelial cells under the influence of the GAM secretome. Therefore, freshly isolated patient-derived tumor endothelial cells were grown as a dense monolayer on culture plate inserts in the presence and absence of GAM-conditioned media. After adding patient-derived autologous T cells, transmigration to the lower chamber was quantified after 24 hours (Suppl. Fig. S3A). In general, we observed a uniform trend toward a decreased transmigration of T cells in the presence of untreated GAM supernatant and the opposite effect upon treatment of GAM with SMI. However, this effect was inconsistent since the level of significance was only reached for some conditions varying among patient samples and for distinct SMI (Suppl. Fig. S3B-D). Instead, the results indicate that GAM secreted molecules with the ability to reduce trans-endothelial T-cell migration into the tumor mass and that GAM treatment can at least partially restore the level of transmigration.

### Impact of SMI-treated GAM on T-cell killing

Since GAMs can impair the ability of intratumoral T cells to kill tumor cells^12,15^, we tested whether CSF1R blockade can improve antitumor immune responses (Fig. 3A). After the generation of pGBM patient-derived tumor cell lines (Fig. 3B), we cocultivated CFSE-stained tumor cells with autologous T cells in the presence of GAM supernatant. After 48 hours, the numbers of live tumor cells (CFSE^+^PI^−^) were quantified. The number of living cells was normalized to tumor cells without T cells and GAM-conditioned medium. While co-incubation of tumor cells with T cells slightly reduced the number of live tumor cells (10.3%), this effect was doubled (20.6%) upon addition of conditioned media from GW2580-treated GAM (p < 0.05) (Fig. 3C-D). By live-cell imaging, we confirmed the induction of T-cell-induced tumor cell apoptosis. Therefore, MitoTracker™-stained tumor cells (red fluorescence) were co-incubated with unstained T cells. Apoptotic tumor cells were identified by the green fluorescent IncuCyte™ Caspase-3/7 Apoptosis Reagent (Suppl. Fig. S4C). Over time, T cells started to accumulate around tumor cells (Suppl. Fig. S4A-B). The highest numbers of apoptotic cells (green) were observed upon incubation with the supernatant of GW2580-treated GAM (Fig. 3E-G).

**Figure 3:**
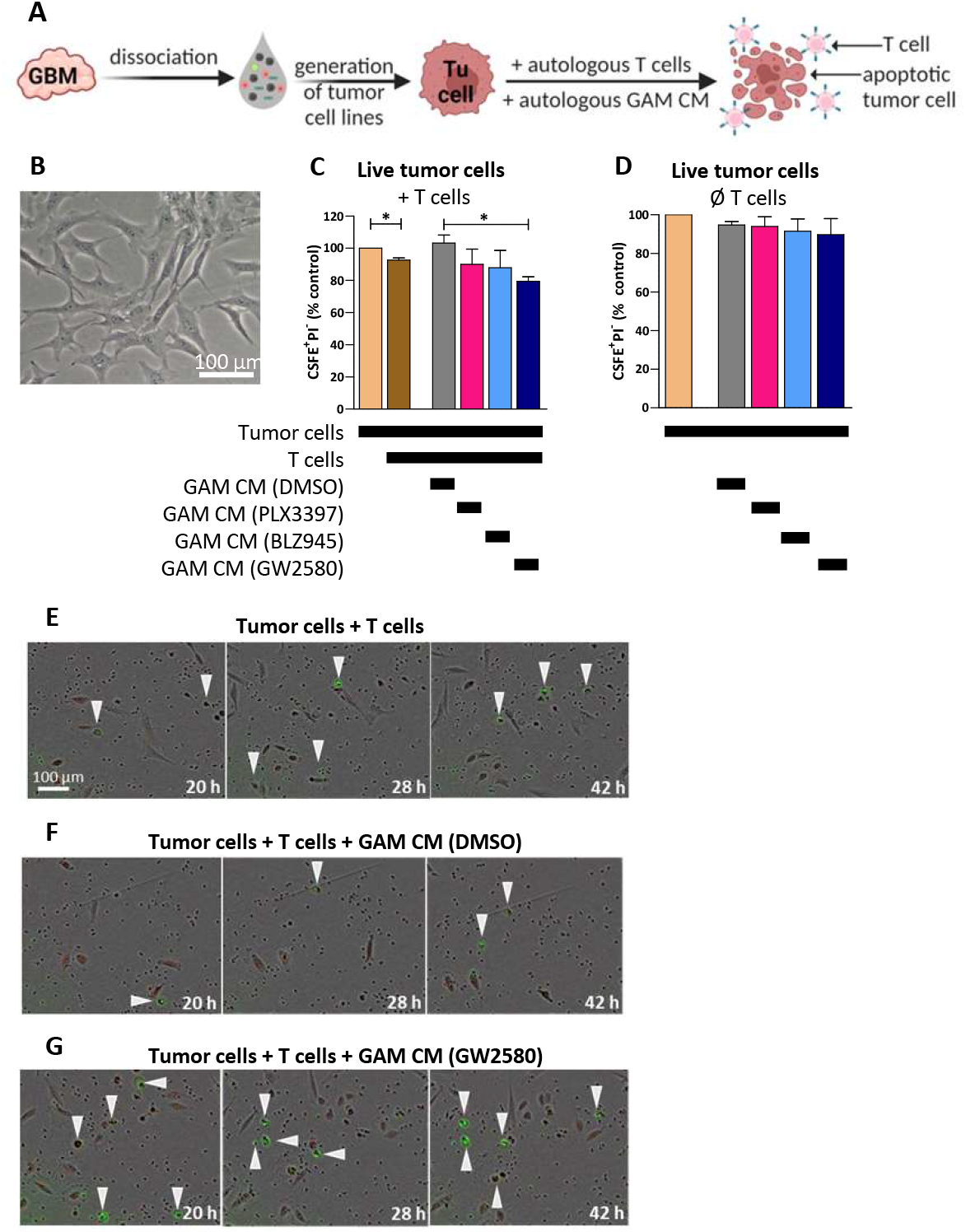
T cells reduce the number of autologous tumor cells in the presence of supernatant of GW2580-treated GAM. **A** pGBM patient-derived tumor cells were incubated with autologous T cells in a 1:10 target-to-effector ratio, and conditioned media of autologous SMI-treated or untreated GAM were added (n = 3). After 48 hours, the cells were collected, stained with PI, and measured by flow cytometry. **B** Microscopic image of a newly generated GBM cell line. **C** Number of CFSE+PI-live tumor cells in the presence of autologous T cells and upon addition of GAM-conditioned media, normalized to the control condition (tumor cells alone). **D** Number of CFSE+PI-live tumor cells in the absence of T cells and upon addition of GAM-conditioned media, normalized to the control condition (no GAM-conditioned media). **E – G** Microscopic images illustrating Caspase-3/7+ apoptotic tumor cells (white arrows) at different time points during the co-cultivation of tumor cells with autologous T cells in the absence of GAM conditioned media (**E**), upon addition of conditioned media of untreated GAM (**F**) and in the presence of conditioned media of GW2580-treated GAM (**G**). Whiskers depict SEM. * p < 0.05; CM = conditioned media; CFSE = carboxyfluorescein succinimidyl ester; PI = propidium iodide; SEM = standard error of the mean.

Altogether, the ability of patient-derived T cells to kill autologous tumor cells through the induction of apoptosis was significantly increased upon treatment with GAM and SMI GW2580.

### CSF1R blockade in pGBM tumor organoids

Finally, the treatment effects of CSF1R blockade were studied in a more natural tumor microenvironment. This was done in pGBM patient-derived tumor organoids, which are believed to recapitulate the composition of the tumor microenvironment as well as intra- and intertumoral heterogeneity more faithfully than in other *in vitro* models^37^. Tumor organoids were generated from freshly resected patient-derived pGBM specimens through bioprinting of single-cell suspensions under nonadherent conditions and then treated with PLX3397, BLZ945, GW2580 or solvent only (0.1% DMSO) (Fig. 4A). Within 5 days, the cells compacted into tumor organoids (Fig. 4B), containing mostly live cells (green) (Fig. 4C) and showing a tissue-like architecture (Fig. 4D). Most of the cells within the tumor organoids consisted of GFAP^+^ tumor cells (Fig. 4G, Suppl. Fig. S5A) in addition to immune cells such as CD68^+^ GAM (Fig. 4E, Suppl. Fig. S5B) and CD3^+^ T cells (Suppl. Fig. S5B). Furthermore, reconstitution of the extracellular matrix (Suppl. Fig. S5A) and the onset of tumor cell proliferation (Fig. 4G, Suppl. Fig. S5A) was observed.

**Figure 4:**
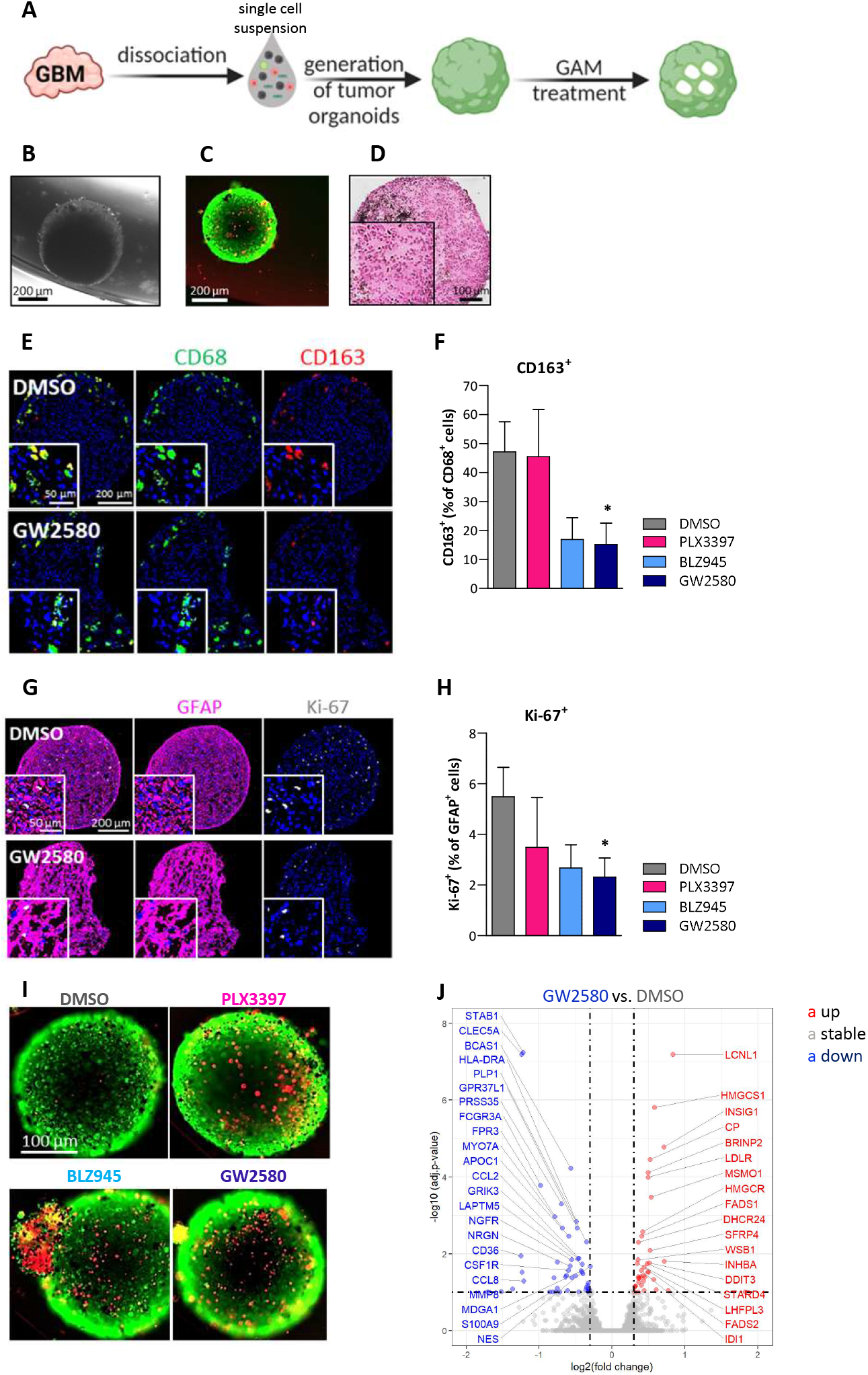
Effective reprogramming and antitumor activity of GW2580-treated GAM in a patient-derived tumor organoid GBM model. Patient-derived GBM organoids were obtained by bioprinting single-cell suspensions. Five-day-old tumor organoids (TO) were treated for 96 hours with PLX3397, BLZ945 or GW2580 (10 μM) compared to the control (0.1% DMSO; n = 4). **B** Bright-field microscopic image of a 5-day-old TO. **C** Live/Dead™ staining of a 5-day-old TO showing mostly green-stained live cells and only a few red-stained dead cells. **D** H&E staining of a 5-day-old TO illustrating a tissue-like structure. **E** Immunofluorescent staining of GW2580-treated and untreated TO for CD68 (green) and CD163 (red) **G** as well as GFAP (pink) and Ki-67 (gray). **F** and **H** show the quantification of CD163+ GAM (out of CD68+) and Ki-67+ proliferating tumor cells (out of GFAP+). Whiskers depict SEM. **I** Live/Dead™ staining of SMI-treated and untreated TO. **J** Volcano plot of differentially expressed genes in GW2580-treated TO compared to untreated TO. Blue = downregulated and red = upregulated genes upon treatment. Level of significance: adj. p value < 0.1, Log2(FC) > 0.3. * p < 0.05; adj. = adjusted; GFAP = glial fibrillary acid protein; SEM = standard error of the mean.

In particular, treatment of pGBM patient-derived tumor organoids with GW2580 not only significantly reduced the percentage of CD68^+^ GAM cells expressing the M2 marker CD163 (Fig. 4E-F) but also the percentage of proliferating Ki-67^+^GFAP^+^ tumor cells (Fig. 4G-H). A trend toward reduced percentages of CD68^+^CD163^+^ and GFAP^+^Ki-67^+^ cells was also seen for BLZ945 and PLX3397 in the same setting (Fig. 4F, H, Suppl. Fig. 6). Interestingly, upon treatment with all three SMIs, LIVE/DEAD cell imaging of tumor organoids revealed an increased portion of dead cells compared to incubation with DMSO alone (Fig. 4I). Moreover, comparative RNA-seq analyses of untreated and treated tumor organoids revealed up to 89 differentially expressed genes (Fig. 4J Suppl. Table 7-8, Suppl. Fig. S7 A-B). Interestingly, GW2580-treated tumor organoids shared several significantly downregulated disease-relevant genes with GW2580-treated GAMs, such as CLEC5A, IL1R2, MARCO, MYO7A, S100A9, and STAB1. CLEC5A has recently been described as an M2 marker and is associated with immunosuppression and poor prognosis in gliomas^38^, while blockade of IL1R2 is able to suppress breast tumorigenesis^39^. MARCO, a well-known scavenger receptor expressed by tumor-associated macrophages, is highly associated with poor prognosis in pancreatic cancer^40^ and was shown to be a mesenchymal protumor marker in GBM^41^. Initial findings on the actin-based motor molecule MYO7A indicate that it plays an important role in the promotion of melanoma progression^42^, while the role of S100A9 as a poor prognosticator in glioma has been previously reported^28^. Finally, STAB1, which is expressed on M2-like macrophages, has been described as a new immunosuppressive molecule in melanoma and seems to be involved in the development of bone metastasis in prostate cancer patients^43,44^. In summary, transcriptomic similarities between treated GAMs and tumor organoids strongly support the reliability of the tumor organoid model and provide the opportunity to also assess transcriptional changes in tumor cells and other stromal cells of the GBM niche. Altogether, even in the more natural tumor microenvironment of tumor organoids where reprogramming of GAMs could be impaired by the presence of tumor cells and other stromal cells, effective proinflammatory and antitumor M1-like activation of patient-derived GAMs was confirmed, especially upon GW2580 treatment.

## Discussion

There is increasing evidence that effective targeting of GAMs by SMIs could have a tremendous impact on the immunosuppressive and tumor-promoting microenvironment of GBM and thus on tumor growth and antitumor immunity^12,19–23,26^. However, which SMI is most beneficial for the treatment of GBM has not been studied thus far. Furthermore, representative data on patient-derived, tumor-educated GAMs are rare because they have often been replaced by macrophages isolated and matured from the peripheral blood of healthy donors, macrophage cell lines or bone-marrow-derived macrophages^22,45–47^. Finally, an understanding of the implications of CSF1R targeting on the adaptive immune system is still lacking. In the present study, we tried to address these limitations by consistently using freshly isolated GAMs, patient-derived tumor cells, tumor-derived endothelial cells and T cells from pGBM patients to study the phenotypical, transcriptional and functional changes of SMI-treated GAMs and their impact on the interaction of tumor and autologous T cells. When comparing the effects of the CSF1R inhibitors PLX3397, BLZ945 and GW2580, the strongest and most significant changes in GAM properties and polarization from an immunosuppressive, tumor-promoting to a proinflammatory, tumor-cytotoxic phenotype were consistently observed for GW2580, followed by less pronounced effects for BLZ945. In contrast, PLX3397 treatment resulted in the weakest changes. Finally, treatment of patient-derived glioblastoma organoids serving as patient avatars by recapitulating the tumor microenvironment and cellular composition of human GBM confirmed a significant reduction in anti-inflammatory GAMs and a downregulation of tumor cell proliferation, particularly in the presence of GW2580.

Our study provides important new insight into the therapeutic potential of different CSF1R-targeting SMIs since PLX3397 and BLZ945 are some of the most broadly studied and clinically most developed SMIs^12,48^. Recently, PLX3397 was approved by the FDA for the treatment of tenosynovial giant cell tumors^49,50^. Although to the best of our knowledge, it is not under clinical investigation, we further included GW2580 in our analysis. Similar to BLZ945, it has a high affinity for CSF1R^51,^ and contrary to PLX3397, in neurodegenerative disease models, it does not lead to unwanted neurotoxic effects since it attenuates microglial proliferation rather than depleting microglial cells^52–54^. In general, CSF1R blockade with all three SMIs was able to reprogram GAMs by i) decreasing the expression of the M2 marker CD163, which is associated with poor survival in GBM^26^, ii) increasing M1-like HLA-DR expression and iii) upregulating M1-related phagocytic ability. However, for PLX3397, the effects were only a trend and failed to reach the level of significance in all analyses, while GW2580 induced the strongest effects. Only the concentration of nitrite in the supernatant of treated GAMs, as a surrogate for the increased M1-like iNOS activation^55^, was significantly increased upon treatment with all three inhibitors but again most effectively for GW2580. This is of particular importance since iNOS is involved in GAM-related cytotoxicity by metabolizing arginine to produce NO and citrulline^56^. Substantial downregulation of M2-like GAMs together with impaired recruitment of GAMs has been observed for PLX3397 and BLZ945 in several preclinical models of murine glioma and xenotransplanted human cell lines^20,22,23^. Furthermore, reduced tumor growth, especially in combination with irradiation, and improved survival were reported for both SMIs^20,22,23,57^. For GW2580, pretreatment of mice before implantation of U251 glioma cells resulted in a decreased number of intratumoral myeloid cells and improved tumor control^58^. However, how GW2580 affects the growth of an already established glioma is not known thus far, but there are promising data from other xenotransplantation models, such as melanoma, hepatocellular carcinoma, and non-small cell lung cancer, describing reduced tumor growth and downregulation of anti-inflammatory M2-like tumor-associated macrophages upon treatment with GW2580^59–61^. Clinically, in addition to many other tumor types, PLX3397 has been administered in a phase II trial to patients suffering from recurrent GBM and in a phase Ib/II trial to newly diagnosed GBM patients combined with radiochemotherapy. In line with our preclinical data, no efficacy was observed for the treatment of recurrent GBM with PLX3397^62^, whereas final results in newly diagnosed GBM have not been published thus far (NCT01790503). BLZ945 is currently under investigation in one active phase I/II trial on advanced solid tumors (NCT02829723) but not in glioma patients, and a final evaluation of patient outcome and tumor control remains to be seen. The weak treatment effects of PLX3397 seen by us and the lack of efficacy in GBM patients are in stark contrast to the promising preclinical data^20^. This might indicate that testing of CSF1R-targeting effects on freshly isolated tumor-educated patient-derived GAM or on tumor organoids might predict response in patients more accurately. In summary, our findings strongly suggest a further preclinical and clinical investigation of GW2580 for the treatment of GBM.

Notably, transcriptome analyses performed on GAMs and tumor organoids substantiated effective reprogramming upon GW2580 treatment. In addition to the downregulation of CD163, MRC1, MMP9, and S100A9 expression, which was also observed upon treatment with PLX3397 and BLZ945, treatment with GW2580 led to the downregulation of some other interesting genes, such as SPP1, which is a glycoprotein that binds to integrins and to CD44 and thereby enhances invasion and metastasis in various cancers, including gliomas^36,63^. Furthermore, significant downregulation of CCL13, which has been indicated as the most informative marker for the M2-like macrophage population in GBM^35^, transforming growth factor beta induced (TGFβI), a marker of the alternative M2b activation status of macrophages^64^, and CD38 was observed upon GAM treatment with GW2580. Interestingly, CD38 upregulation is known to mediate suppression of cytotoxic T cells via adenosine receptor signaling and has been associated with functional impairment of cytotoxic T cells in animal models of lung cancer and melanomas acquiring resistance to anti-PD-1 and anti-PD-L1 therapies^34^. In addition, CD38^hi^ myeloid-derived suppressor cells (MDSCs) seem to be more effective in suppressing activated T cells and promoting tumor growth than CD38^lo^ MDSCs^65^. Accordingly, downregulation of CD38 gene expression through GW2580 may sustain T-cell-mediated antitumor effects by impairing GAM immunosuppressive functions. Moreover, gene set enrichment analyses and cytokine quantification revealed significantly lower levels of transcripts related to IL10 signaling and the IL-6-STAT3 signaling pathway in GW2580-treated GAMs along with reduced IL-6 levels in their supernatant. This observation is crucial since IL-6 regulates several hallmarks of cancer. High levels of IL-6 in the serum and tumor tissues of several cancer patients^32^ are associated with aggressive tumor growth (34) and shorter survival^66^.

Based on our findings and previous observations by others that myeloid cells are key regulators of the immunosuppressive tumor microenvironment by decreasing iNOS activity^15,67^ and suppressing T-cell function^17,18,18,68^, an important aim of the present study was to determine whether reprogramming GAMs by CSF1R blockade could improve antitumor T-cell responses. To this end, we tested the ability of patient-derived T cells to transmigrate through a layer of tumor-derived endothelial cells and to control the growth of tumor cells in the presence of untreated and treated GAM supernatant. While we observed a partially significant but inconsistent improvement in T-cell transmigration after treatment with all three SMIs, regarding the impact on tumor cell growth, GW2580 outperformed all other SMIs by significantly reducing the number of live tumor cells in the presence of T cells. Notably, in the absence of T cells, this killing effect was not observed. Furthermore, live-cell imaging of tumor-T-cell cocultures confirmed the increased ability of T cells to induce tumor cell apoptosis under the influence of supernatant from GW2580-treated GAMs. The beneficial effects observed for GW2580 treatment strongly support further preclinical and clinical studies to elucidate whether additional treatment with this SMI in an immunotherapeutic setting could improve the efficacy of antitumor T-cell responses in GBM. This is supported by previous data in a pancreatic cancer mouse model where GW2580 treatment improved the response to immune checkpoint inhibitors^45^. Even for the less effective SMI PLX3397, a combination with a dendritic cell vaccine and anti-PD-1 antibodies showed remarkable effects by improving T-cell infiltration and overall survival of GL261-bearing mice^69^. For the application of BLZ945 together with anti-PD-L1 antibodies, increased infiltration of T cells was observed in an artificial 3D model consisting of GBM cells, immortal macrophages and immortalized brain endothelial cells^47^.

Altogether, the results of our comparative analysis of CSF1R-targeting SMIs and the most effective reprogramming of patient-derived GAMs by GW2580 strongly suggest that additional treatment with GW2580 could be an important pillar in future (immune-)therapeutic settings for the treatment of GBM. Future studies will need to show if these promising effects can be translated into an effective clinical use.

## Acknowledgment

We thank Ronja Trunk, Oskar Fürst, Lisa Petermann, and Jan Moser for skillful technical assistance.

