## supplemental figures for "Reprogramming M2-polarized patient-derived glioblastoma associated microglia/macrophages via CSF1R inhibition"

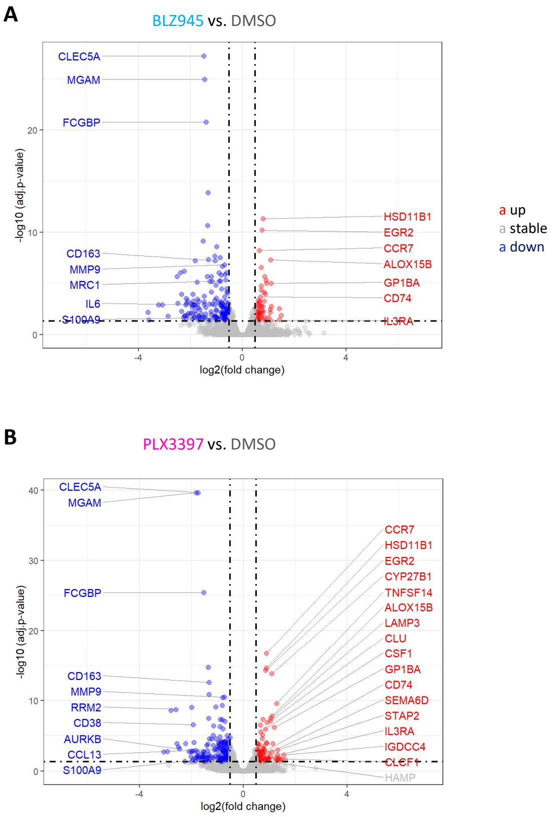


**Supplementary figure 1: Differential gene expression of BLZ945- and PLX3397-treated GAM.**

Volcano plots of differentially expressed genes in BLZ945- (**A)** and PLX3397-treated (**B)** compared to untreated GAM. Blue = downregulated and red = upregulated genes. Level of significance: adj. p value < 0.05, $Log_2(FC) > 0.5$. Log = logarithm; FC = fold change; adj. = adjusted.

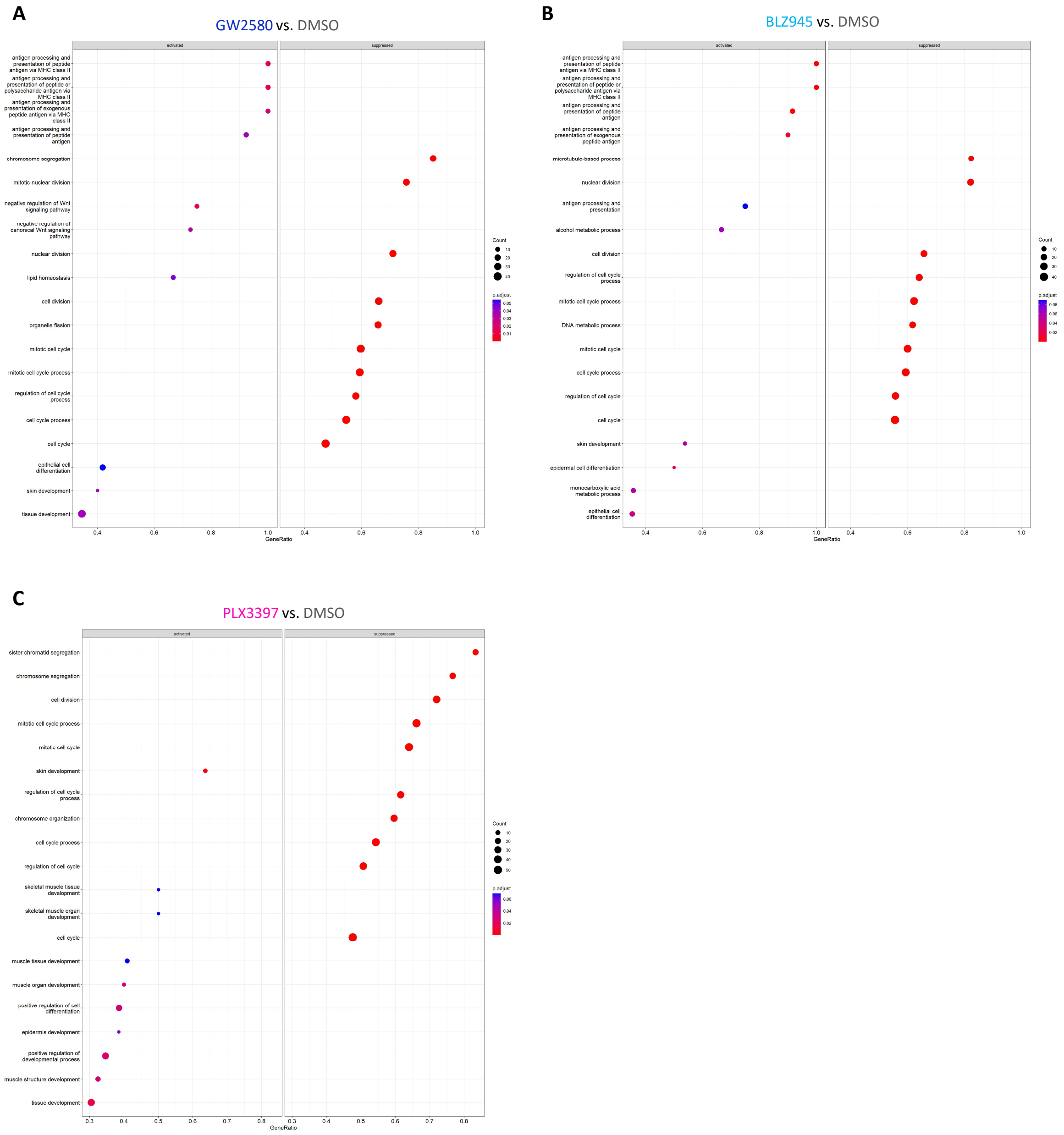


**Supplementary figure 2: Gene set enrichment analyses of RNA sequencing data identifies enhanced antigen presentation in GW2580 and BLZ945-treated GAM.**

The 20 top differentially expressed GO terms in GW2580-, BLZ945-, and PLX3397-treated GAM as compared to untreated GAM are shown in **A**, **B**, and **C**, respectively. Each dot represents one gene set, and the size of each dot depends on the number of genes that are contained in the set. A color code is used to represent the adjusted p value. p adjust = adjusted p-value.

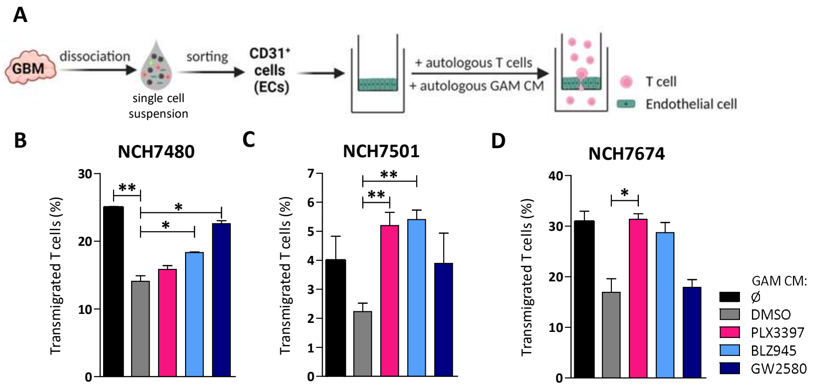


**Supplementary figure 3: Increased transmigration of patient-derived T cells through a dense blood-brain-barrier-like autologous endothelial cell layer upon treatment of GAM**

**A** pGBM patient-derived endothelial cells forming a dense monolayer were exposed to conditioned media of SMI-treated or untreated GAM and the number of autologous T cells transmigrating through the tumor-derived endothelial cell layer was quantified (n = 3 done in triplicates). Endothelial cells cultivated with medium alone were used as a control. **B-D** Percentage of transmigrated T cells as quantified by flow cytometry. Whiskers depict SEM. * $p < 0.05$; ** $p < 0.01$.; ECs = endothelial cells; CM = conditioned media; SEM = standard error of the mean.

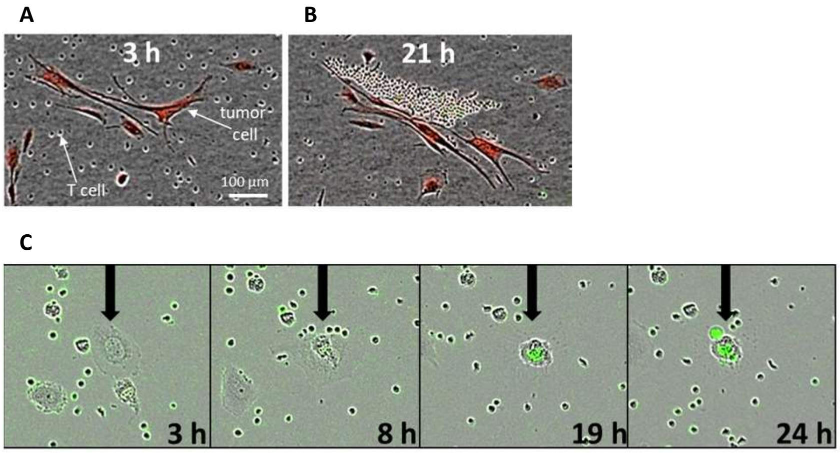


**Supplementary figure 4: Live-cell imaging demonstrates accumulation of autologous T cells around tumor cells and the induction of tumor cell apoptosis.**

pGBM patient-derived tumor cells were incubated with autologous T cells in a 1:10 target-to-effector ratio and exposed to conditioned media of SMI-treated or untreated GAM. Tumor cells were stained with the red fluorescent dye MytoTracker$^{TM}$ to distinguish them from unstained T cells. The green fluorescent IncuCyte$^{TM}$ Caspase-3/7 Apoptosis Regent was added to identify apoptotic cells. Live-cell imaging was performed for 48 hours using the IncuCyte$^{TM}$ Zoom Instrument. **A** During the first hours of co-cultivation of T cells with autologous tumor cells, T cells (small, round cells) showed a uniform distribution within the well. Thereafter, T cells accumulated around tumor cells (elongated, red-stained cells). **B** Example of a tumor cell (black arrow) undergoing apoptosis and turning green because of the presence of the green fluorescent IncuCyte$^{TM}$ Caspase-3/7 Apoptosis Reagent.

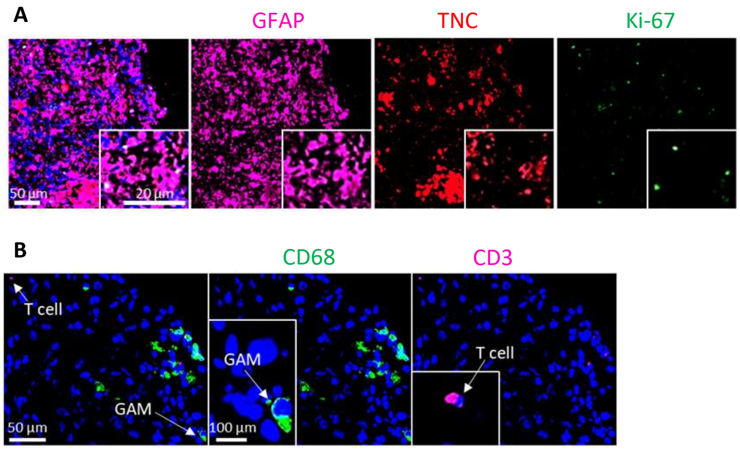


**Supplementary figure 5: pGBM patient-derived tumor organoids maintain cellular composition, extracellular matrix, and proliferative activity of the parental tumor.**

**A** Immunofluorescent staining of a 9 days-old tumor organoid containing GFAP$^+$ tumor cells (pink) that retain the ability to produce extracellular matrix (TNC$^+$ staining in red) and to proliferate (Ki-67$^+$ staining in green). **B** Immunofluorescent stainings of a 9 days-old tumor organoid with antibodies against CD68 (green) and CD3 (pink). GFAP = glial fibrillary acid protein; TNC = tenascin C; Ki-67 = marker of proliferation Ki-67; GAM = glioblastoma-associated microglia/macrophages.

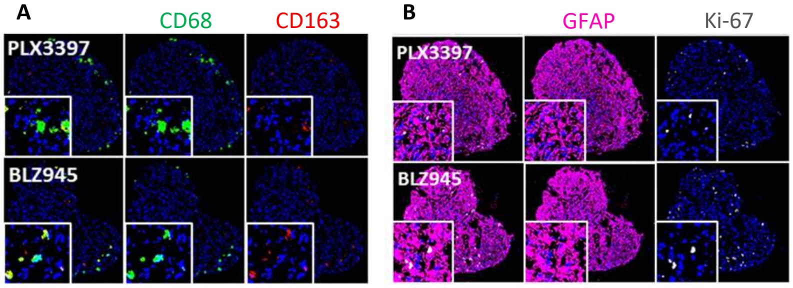


**Supplementary figure 6: Changes in the cellular composition and proliferative activity in pGBM patient-derived tumor organoids upon treatment with PLX3397 or BLZ945.**

**A** Immunofluorescent staining of SMI-treated tumor organoids with antibodies against CD68 (green) and CD163 (red). **B** and GFAP (pink) and Ki-67 (grey). GFAP = glial fibrillary acid protein; Ki-67 = marker of proliferation Ki-67.

**A**

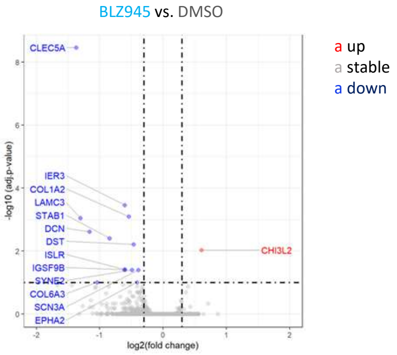


**B**

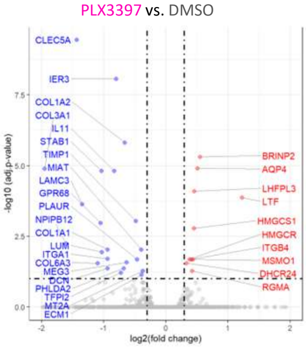


**Supplementary figure 7: Gene expression profile of BLZ945- and PLX3397-treated pGBM patient-derived organoids.**
Volcano plots of differentially expressed genes in BLZ945- (**A)** and PLX3397-treated (**B)** tumor organoids compared to untreated organoids. Blue = downregulated and red = upregulated genes upon GAM treatment with the SMI. Level of significance: adj. p value < 0.1, $Log_2$(FC) > 0.3. Log = logarithm; FC = fold change; adj. = adjusted.
