## supplemental tables for "Reprogramming M2-polarized patient-derived glioblastoma associated microglia/macrophages via CSF1R inhibition"

**Supplementary table 1: Clinical overview of patients´ sample**

| Sample ID | Age at diagnosis (years) | Sex | IDH1-mutation | MGMT-methylation | Subtype | GAM Phenotyping | Tumor organoids |
|---|---|---|---|---|---|---|---|
| NCH6723 | 63 | m | wild type | hypermethylated | N/A | x | |
| NCH6816 | 76 | m | wild type | non-hypermethylated | N/A | x | |
| NCH6835 | 70 | m | wild type | non-hypermethylated | mesenchymal | x | |
| NCH6866 | 56 | m | wild type | non-hypermethylated | mesenchymal | x | |
| NCH7025 | 63 | m | wild type | hypermethylated | N/A | x | |
| NCH7065 | 78 | m | wild type | non-hypermethylated | N/A | x | |
| NCH7163 | 67 | f | wild type | non-hypermethylated | mesenchymal | x | |
| NCH7460 | 76 | f | wild type | hypermethylated | middle line | x | |
| NCH7480 | 55 | m | wild type | non-hypermethylated | RTK II | x | |
| NCH7501 | 73 | f | wild type | non-hypermethylated | RTK II | x | |
| NCH7507 | 59 | f | wild type | hypermethylated | mesenchymal | x | |
| NCH7583 | 73 | m | wild type | hypermethylated | middle line | x | |
| NCH7674 | 60 | m | wild type | non-hypermethylated | mesenchymal | x | |
| NCH7713 | 79 | m | wild type | non-hypermethylated | N/A | x | |
| NCH7925 | 69 | m | wild type | hypermethylated | N/A | x | |
| NCH7952 | 59 | f | wild type | non-hypermethylated | RTK II | x | |
| NCH8110 | 71 | f | wild type | non-hypermethylated | RTK I | | x |
| NCH8245 | 73 | f | wild type | non-hypermethylated | N/A | x | |
| NCH8261 | 66 | m | wild type | hypermethylated | N/A | x | |
| NCH8442 | 51 | m | wild type | non-hypermethylated | N/A | | x |
| NCH8476 | 81 | m | wild type | hypermethylated | N/A | | x |
| NCH9103 | 66 | m | wild type | hypermethylated | N/A | | x |

| Characteristics | | |
|---|---|---|
| **Age (years)** | Median ± SD 67.4 (8.4) | Range (51 - 81) |
| **Sex** | **Frequency** | **Percentage** |
| male | 15 | 68.2 |
| female | 7 | 31.8 |

Abbreviations: IDH1 = isocitrate dehydrogenase (NADP(+))1; MGMT = O-6-methylguanine-DNA methyltransferase; GAM = glioblastoma-associated microglia/macrophages; m = male; f = female

**Supplementary table 2: Antibodies for flow cytometry**

| Specificity | Species | Clone | isotype | Fluorophore | Dilution (concentration) | Manufacturer | Order number |
|---|---|---|---|---|---|---|---|
| **CD68** | mouse | Ki-M7 | IgG1 | FITC | 1:5 (2.6 µg/ml) | Caltag. Buckingham. UK | GM4152 |
| **CD163** | mouse | GHI/61 | IgG1.k | PE | 1:20 (10 µg/ml) | BD Biosciences. Heidelberg. DE | 330636 |
| **HLA-DR** | mouse | G46-6 | IgG2b.k | PE-Cy7 | 1:20 (5 µg/ml) | BD Biosciences. Heidelberg. DE | 560651 |

Abbreviations: CD = cluster of differentiation; HLA-DR = major histocompatibility complex. class II. DR

**Supplementary table 3: Primary antibodies for immunofluorescent stainings**

| Specificity | Species | Clone | isotype | Dilution (concentration) | Manufacturer | Order number |
|---|---|---|---|---|---|---|
| **GFAP** | rabbit | polyclonal | IgG1.k | 1:600 (4.7 µg/ml) | DAKO. Hamburg. DE | Z0334 |
| **TNC** | mouse | BC-24 | IgG1 | 1:80 (1.9 µg/ml) | Sigma-Aldrich. Taufkirchen. DE | T2551 |
| **Ki-67** | mouse | B56 | IgG1.k | 1:25 (2 µg/ml) | BD Bioscience. Bedford. US | MKI67 |
| **CD68** | mouse | EBM11 | IgG1.k | 1:25 (10.9 µg/ml) | DAKO. Hamburg. DE | M0718 |
| **CD163** | mouse | EDHu-1 | IgG1 | 1:300 (3.34 µg/ml) | Bio-Rad Laboratories. Hercules. USA | MCA1853 |
| **CD3** | rabbit | polyclonal | IgG1 | 1:100 (50 µg/ml) | DAKO. Hamburg. DE | A0452 |

Abbreviations: GFAP = glial fibrillary acid protein; TNC = tenascin C; Ki-67 = marker of proliferation Ki-67; CD = cluster of differentiation

**Supplementary table 4: Fluorophore-conjugated secondary antibodies**

| Specificity | Species | Fluorophore | Dilution (concentration) | Manufacturer | Order number |
|---|---|---|---|---|---|
| **mouse IgG (H+L)** | chicken | AlexaFluor® 647 | 1:200 (10 µg/ml) | Thermo Fisher Scientific. Walthan. USA | A-21463 |
| **rabbit IgG (H+L)** | goat | AlexaFluor® 647 | 1:200 (10 µg/ml) | Thermo Fisher Scientific. Walthan. USA | A-21245 |
| **mouse IgG (H+L)** | goat | AlexaFluor® 555 | 1:400 (5 µg/ml) | Thermo Fisher Scientific. Walthan. USA | A-21424 |
| **Zenon IgG1 Labeling Kit** | mouse | AlexaFluor® 488 | | Thermo Fisher Scientific. Walthan. USA | Z-25002 |

### Supplementary table 5A: Common differentially expressed genes upon CSF1R blockade in pGBM patient-derived GAM (downregulated)

| | GW2580 | | BLZ945 | | PLX3397 | |
|---|---|---|---|---|---|---|
| Gene | Log(FC) | Adj. p-value | Log(FC) | Adj. p-value | Log(FC) | Adj. p-value |
| CCL7 | -3.15 | 1.47E-03 | -3.23 | 1.22E-03 | -3.06 | 2.01E-03 |
| RUFY4 | -3.06 | 1.23E-03 | -3.07 | 1.26E-03 | -2.92 | 1.74E-03 |
| UBE2C | -2.47 | 7.81E-04 | -2.48 | 8.61E-04 | -2.02 | 8.47E-03 |
| IL1R2 | -2.42 | 6.09E-07 | -2.38 | 9.40E-07 | -2.79 | 2.27E-09 |
| RRM2 | -2.31 | 2.42E-07 | -2.24 | 6.44E-07 | -2.59 | 1.77E-09 |
| DTL | -2.28 | 9.46E-05 | -2.11 | 4.95E-04 | -2.22 | 1.33E-04 |
| CDK1 | -2.11 | 1.82E-03 | -1.68 | 2.73E-02 | -1.80 | 1.12E-02 |
| FCN1 | -2.10 | 2.77E-10 | -1.82 | 6.00E-08 | -2.00 | 8.57E-10 |
| AURKB | -2.07 | 6.13E-03 | -1.85 | 2.17E-02 | -2.45 | 7.19E-04 |
| MMP8 | -2.06 | 2.91E-03 | -1.93 | 8.67E-03 | -1.75 | 2.01E-02 |
| TROAP | -1.94 | 2.65E-02 | -2.33 | 5.32E-03 | -1.84 | 3.78E-02 |
| SHCBP1 | -1.93 | 2.42E-03 | -2.33 | 1.15E-04 | -1.98 | 1.47E-03 |
| PLK1 | -1.92 | 8.31E-05 | -1.90 | 1.14E-04 | -1.90 | 9.22E-05 |
| E2F2 | -1.87 | 1.12E-02 | -1.99 | 7.51E-03 | -2.56 | 1.33E-04 |
| MT1HL1 | -1.86 | 2.35E-03 | -1.81 | 4.44E-03 | -1.47 | 3.27E-02 |
| CDC45 | -1.83 | 9.23E-03 | -2.18 | 1.27E-03 | -2.10 | 1.63E-03 |
| CD38 | -1.82 | 2.48E-06 | -1.74 | 7.37E-06 | -1.93 | 2.65E-07 |
| MT1H | -1.78 | 1.64E-03 | -1.83 | 1.27E-03 | -1.32 | 4.72E-02 |
| MYBL2 | -1.74 | 2.60E-03 | -2.51 | 2.11E-06 | -1.75 | 2.04E-03 |
| CDC20 | -1.74 | 3.77E-03 | -1.92 | 1.19E-03 | -1.59 | 1.05E-02 |
| CCL2 | -1.73 | 9.49E-03 | -1.67 | 1.82E-02 | -1.61 | 2.10E-02 |
| MT1G | -1.71 | 3.04E-04 | -1.73 | 3.52E-04 | -1.37 | 9.13E-03 |
| MT1A | -1.65 | 3.03E-03 | -1.88 | 5.51E-04 | -1.40 | 1.73E-02 |
| PID1 | -1.65 | 3.03E-03 | -1.70 | 2.41E-03 | -1.49 | 1.08E-02 |
| MGAM | -1.65 | 1.54E-32 | -1.46 | 1.04E-25 | -1.82 | 2.35E-40 |
| TYMS | -1.62 | 6.57E-03 | -1.85 | 1.27E-03 | -1.83 | 1.19E-03 |
| MT1L | -1.61 | 1.15E-02 | -1.99 | 9.51E-04 | -1.39 | 4.17E-02 |
| AC073072.5 | -1.58 | 1.23E-03 | -1.67 | 6.07E-04 | -1.53 | 1.63E-03 |
| GPR82 | -1.58 | 1.19E-04 | -1.18 | 1.16E-02 | -1.37 | 1.27E-03 |
| ETV5 | -1.57 | 1.18E-02 | -1.52 | 2.12E-02 | -1.58 | 1.14E-02 |
| CLEC5A | -1.57 | 4.79E-32 | -1.48 | 5.35E-28 | -1.75 | 2.35E-40 |
| CEP55 | -1.54 | 3.57E-02 | -2.15 | 8.61E-04 | -2.01 | 1.66E-03 |
| CDT1 | -1.51 | 1.31E-02 | -1.59 | 9.17E-03 | -1.40 | 2.57E-02 |
| CCNA2 | -1.49 | 9.23E-03 | -1.64 | 3.56E-03 | -1.59 | 4.19E-03 |
| FPR2 | -1.49 | 2.82E-05 | -1.25 | 1.05E-03 | -1.37 | 1.53E-04 |
| MT1P3 | -1.49 | 1.03E-09 | -1.50 | 6.96E-10 | -1.15 | 8.69E-06 |
| GNG2 | -1.47 | 1.94E-07 | -1.35 | 2.03E-06 | -1.22 | 2.79E-05 |
| FCGBP | -1.45 | 2.51E-23 | -1.40 | 1.68E-21 | -1.53 | 3.86E-26 |
| PRC1 | -1.44 | 1.71E-02 | -1.44 | 2.03E-02 | -1.54 | 8.15E-03 |
| MT1M | -1.43 | 7.42E-04 | -1.46 | 6.09E-04 | -1.15 | 1.32E-02 |
| IRG1 | -1.41 | 2.77E-06 | -1.00 | 3.63E-03 | -1.16 | 2.38E-04 |
| FOXM1 | -1.41 | 1.11E-02 | -1.26 | 4.06E-02 | -1.52 | 4.35E-03 |
| IL6 | -1.39 | 1.79E-02 | -1.74 | 1.24E-03 | -1.43 | 1.33E-02 |
| CCL13 | -1.38 | 3.48E-06 | -1.06 | 1.27E-03 | -1.07 | 8.11E-04 |
| FPR1 | -1.38 | 3.00E-12 | -1.33 | 2.25E-11 | -1.33 | 1.28E-11 |
| VSIG4 | -1.34 | 4.09E-15 | -1.32 | 1.32E-14 | -1.35 | 1.82E-15 |
| MT1X | -1.25 | 2.70E-07 | -1.31 | 4.73E-08 | -1.06 | 2.58E-05 |
| GPR34 | -1.24 | 9.14E-05 | -1.00 | 5.09E-03 | -1.31 | 2.58E-05 |
| CD300E | -1.22 | 1.75E-03 | -1.06 | 1.45E-02 | -1.09 | 7.85E-03 |
| EDN1 | -1.18 | 7.45E-05 | -1.19 | 7.00E-05 | -0.75 | 3.94E-02 |
| CD163 | -1.15 | 1.03E-09 | -1.05 | 6.00E-08 | -1.32 | 2.67E-13 |
| AC060834.2 | -1.13 | 5.16E-03 | -1.02 | 1.93E-02 | -0.91 | 4.35E-02 |
| FLT1 | -1.12 | 1.64E-05 | -1.17 | 6.21E-06 | -0.98 | 3.16E-04 |
| TREM1 | -1.11 | 1.32E-04 | -1.06 | 4.61E-04 | -0.82 | 1.34E-02 |
| MT1E | -1.06 | 5.82E-05 | -1.00 | 2.63E-04 | -0.73 | 1.79E-02 |
| RNASE2 | -1.05 | 8.62E-04 | -0.83 | 2.38E-02 | -0.86 | 1.20E-02 |
| MAP3K7CL | -1.04 | 9.23E-03 | -1.00 | 1.78E-02 | -0.95 | 2.45E-02 |
| SLC9A7P1 | -1.03 | 4.71E-04 | -1.25 | 6.27E-06 | -0.99 | 8.92E-04 |
| THBD | -1.02 | 1.01E-04 | -1.19 | 2.11E-06 | -1.34 | 3.26E-08 |
| WFDC21P | -1.02 | 4.94E-05 | -0.91 | 6.94E-04 | -0.72 | 1.28E-02 |
| P3H2 | -1.01 | 1.41E-03 | -0.83 | 2.05E-02 | -0.83 | 1.45E-02 |
| MRC1 | -0.99 | 1.67E-07 | -0.88 | 5.98E-06 | -0.74 | 2.75E-04 |
| GIMAP6 | -0.98 | 3.36E-04 | -0.82 | 6.68E-03 | -0.75 | 1.37E-02 |
| MT2A | -0.97 | 4.03E-07 | -1.06 | 2.05E-08 | -0.80 | 7.75E-05 |
| IL1B | -0.96 | 7.72E-09 | -0.99 | 2.56E-09 | -0.72 | 6.73E-05 |
| PTX3 | -0.96 | 9.19E-03 | -1.14 | 1.05E-03 | -1.12 | 1.11E-03 |
| S100A9 | -0.94 | 2.57E-02 | -0.98 | 2.17E-02 | -1.00 | 1.34E-02 |
| HRH2 | -0.94 | 8.91E-05 | -0.81 | 1.54E-03 | -1.14 | 3.66E-07 |
| MT1F | -0.91 | 2.01E-04 | -1.02 | 1.53E-05 | -0.70 | 1.12E-02 |
| S100B | -0.91 | 7.04E-05 | -0.83 | 5.82E-04 | -0.65 | 1.27E-02 |
| NLRP3 | -0.90 | 5.24E-05 | -0.75 | 2.15E-03 | -0.83 | 3.07E-04 |
| SPP1 | -0.90 | 3.57E-02 | -0.94 | 2.80E-02 | -0.90 | 3.45E-02 |
| TNFRSF11A | -0.89 | 1.05E-03 | -0.76 | 1.06E-02 | -0.95 | 3.07E-04 |
| FCAR | -0.89 | 4.16E-04 | -0.85 | 1.11E-03 | -0.96 | 9.02E-05 |
| CDK6 | -0.84 | 9.05E-06 | -0.80 | 3.59E-05 | -0.75 | 1.40E-04 |
| STAB1 | -0.82 | 9.79E-06 | -0.61 | 5.03E-03 | -1.06 | 7.48E-10 |
| CORO1A | -0.81 | 1.64E-05 | -0.84 | 5.98E-06 | -0.90 | 4.61E-07 |
| SERPINB9 | -0.80 | 2.26E-04 | -0.63 | 1.10E-02 | -0.53 | 4.67E-02 |
| PROCR | -0.80 | 5.43E-04 | -0.79 | 8.00E-04 | -0.75 | 1.46E-03 |
| RP11-1036E20.7 | -0.78 | 3.57E-03 | -0.64 | 3.95E-02 | -0.80 | 2.27E-03 |
| CXCL3 | -0.76 | 2.04E-07 | -0.76 | 2.15E-07 | -0.61 | 1.23E-04 |
| TMPO | -0.76 | 2.69E-03 | -0.78 | 2.04E-03 | -0.74 | 3.36E-03 |
| GLIS3 | -0.72 | 1.46E-02 | -0.77 | 9.35E-03 | -0.64 | 4.39E-02 |
| VNN1 | -0.71 | 7.04E-05 | -0.65 | 5.68E-04 | -0.72 | 6.26E-05 |
| CCND1 | -0.70 | 1.06E-05 | -0.85 | 2.89E-08 | -0.62 | 2.36E-04 |
| LMO2 | -0.70 | 3.22E-03 | -0.76 | 1.06E-03 | -0.88 | 5.15E-05 |
| C1orf21 | -0.70 | 5.88E-03 | -0.69 | 8.30E-03 | -0.67 | 8.65E-03 |
| TIMP1 | -0.69 | 8.91E-05 | -0.81 | 1.56E-06 | -0.89 | 4.46E-08 |
| DUSP6 | -0.68 | 1.01E-04 | -0.54 | 7.84E-03 | -0.70 | 7.73E-05 |
| FCGR1A | -0.67 | 9.23E-03 | -0.69 | 9.91E-03 | -0.88 | 1.23E-04 |
| MMP9 | -0.67 | 1.67E-07 | -0.67 | 1.49E-07 | -0.80 | 3.43E-11 |
| TMPO-AS1 | -0.67 | 9.23E-03 | -0.64 | 1.76E-02 | -0.77 | 1.29E-03 |
| RP11-465L10.10 | -0.67 | 1.39E-06 | -0.67 | 1.40E-06 | -0.81 | 4.59E-10 |
| MARCO | -0.66 | 5.30E-03 | -0.70 | 2.70E-03 | -0.74 | 9.77E-04 |
| NT5DC2 | -0.64 | 4.16E-04 | -0.58 | 3.37E-03 | -0.69 | 1.19E-04 |
| MYC | -0.64 | 1.21E-03 | -0.61 | 2.92E-03 | -0.62 | 1.85E-03 |
| LRRC25 | -0.64 | 4.19E-06 | -0.64 | 4.53E-06 | -0.71 | 8.16E-08 |
| SLA | -0.64 | 1.15E-03 | -0.54 | 1.52E-02 | -0.62 | 1.50E-03 |
| FCGR1B | -0.64 | 3.66E-02 | -0.67 | 2.97E-02 | -0.87 | 8.37E-04 |
| CPAMD8 | -0.64 | 3.80E-02 | -0.64 | 4.56E-02 | -0.63 | 3.78E-02 |
| HNMT | -0.63 | 3.22E-03 | -0.60 | 8.17E-03 | -0.66 | 1.63E-03 |
| GPR84 | -0.63 | 8.62E-04 | -0.59 | 2.92E-03 | -0.54 | 7.81E-03 |
| APBB3 | -0.63 | 2.35E-03 | -0.57 | 1.03E-02 | -0.64 | 1.47E-03 |
| TMEM106A | -0.62 | 2.63E-05 | -0.53 | 8.22E-04 | -0.59 | 9.05E-05 |
| FCGR1C | -0.62 | 3.99E-02 | -0.74 | 7.71E-03 | -0.92 | 1.47E-04 |
| KMO | -0.60 | 3.00E-03 | -0.51 | 2.52E-02 | -0.52 | 1.60E-02 |
| FZD1 | -0.59 | 4.02E-04 | -0.51 | 4.99E-03 | -0.81 | 5.07E-08 |
| ARHGAP4 | -0.57 | 7.45E-05 | -0.60 | 3.36E-05 | -0.62 | 9.49E-06 |
| CKAP4 | -0.57 | 3.57E-03 | -0.69 | 2.02E-04 | -0.63 | 7.71E-04 |
| P2RY8 | -0.55 | 2.92E-02 | -0.71 | 1.69E-03 | -0.89 | 1.41E-05 |
| EMILIN2 | -0.55 | 3.90E-03 | -0.53 | 8.17E-03 | -0.54 | 4.85E-03 |
| ADA | -0.54 | 3.80E-04 | -0.60 | 6.70E-05 | -0.56 | 1.62E-04 |
| EHD1 | -0.52 | 3.46E-04 | -0.56 | 8.89E-05 | -0.51 | 5.10E-04 |
| LAMC1 | -0.51 | 2.77E-02 | -0.53 | 2.25E-02 | -0.55 | 1.28E-02 |

Abbreviations: Log = logarithm; FC = fold change; adj. = adjusted

**Supplementary table 5B: Common differentially expressed genes upon CSF1R blockade in pGBM patient-derived GAM (upregulated)**

| | GW2580 | | BLZ945 | | PLX3397 | |
|---|---|---|---|---|---|---|
| Gene | Log(FC) | Adj. p-value | Log(FC) | Adj. p-value | Log(FC) | Adj. p-value |
| SCIN | 0.56 | 1.91E-03 | 0.58 | 1.10E-03 | 0.58 | 1.02E-03 |
| CYP27B1 | 0.56 | 1.83E-03 | 0.62 | 4.62E-04 | 1.10 | 1.55E-14 |
| CD22 | 0.57 | 1.41E-02 | 0.60 | 9.16E-03 | 0.63 | 3.30E-03 |
| KLHL6 | 0.57 | 2.40E-04 | 0.50 | 2.66E-03 | 0.58 | 1.41E-04 |
| CCR7 | 0.58 | 6.39E-07 | 0.66 | 6.13E-09 | 0.89 | 1.93E-17 |
| TSKU | 0.65 | 7.05E-04 | 0.57 | 6.89E-03 | 0.55 | 8.05E-03 |
| CD74 | 0.68 | 3.35E-04 | 0.70 | 2.16E-04 | 0.57 | 5.34E-03 |
| EGR2 | 0.72 | 1.03E-09 | 0.76 | 6.26E-11 | 0.87 | 4.97E-15 |
| HSD11B1 | 0.73 | 1.03E-09 | 0.81 | 4.84E-12 | 0.89 | 2.29E-15 |
| LAMP3 | 0.74 | 7.26E-04 | 0.92 | 5.11E-06 | 1.06 | 3.26E-08 |
| C10orf105 | 0.75 | 1.41E-02 | 0.82 | 5.37E-03 | 0.70 | 2.63E-02 |
| TCN2 | 0.75 | 3.70E-05 | 0.66 | 7.75E-04 | 0.54 | 9.35E-03 |
| CES1 | 0.77 | 3.73E-06 | 0.73 | 1.78E-05 | 0.75 | 7.86E-06 |
| ACSM5 | 0.79 | 1.47E-03 | 0.78 | 1.81E-03 | 0.68 | 1.10E-02 |
| GCHFR | 0.79 | 4.87E-06 | 0.62 | 1.33E-03 | 0.62 | 1.27E-03 |
| HTRA4 | 0.79 | 1.12E-03 | 0.70 | 9.16E-03 | 0.89 | 1.04E-04 |
| CES1P1 | 0.79 | 3.92E-06 | 0.73 | 4.67E-05 | 0.74 | 2.58E-05 |
| SEMA6D | 0.82 | 2.35E-02 | 0.85 | 1.94E-02 | 0.97 | 2.80E-03 |
| NUPR1 | 0.86 | 3.43E-05 | 0.64 | 8.17E-03 | 0.64 | 6.28E-03 |
| SNAI3 | 0.87 | 2.51E-04 | 0.68 | 1.39E-02 | 0.77 | 1.82E-03 |
| CFD | 0.88 | 7.74E-04 | 0.77 | 7.53E-03 | 0.62 | 4.72E-02 |
| CLU | 0.88 | 3.53E-05 | 0.86 | 7.84E-05 | 1.06 | 7.20E-08 |
| CEBPA | 0.90 | 9.79E-06 | 0.71 | 1.81E-03 | 0.77 | 3.16E-04 |
| COL8A2 | 0.93 | 1.74E-07 | 0.86 | 2.06E-06 | 0.87 | 1.11E-06 |
| ST14 | 0.97 | 3.52E-08 | 0.69 | 6.00E-04 | 0.93 | 1.33E-07 |
| ALOX15B | 0.98 | 2.55E-06 | 1.09 | 4.84E-08 | 1.11 | 1.55E-08 |
| MLPH | 0.99 | 9.23E-03 | 1.13 | 1.81E-03 | 0.94 | 1.64E-02 |
| GPR146 | 1.02 | 9.79E-06 | 0.76 | 5.06E-03 | 0.82 | 1.29E-03 |
| OLFM2 | 1.03 | 5.72E-07 | 0.93 | 1.02E-05 | 0.78 | 6.05E-04 |
| CXCR4 | 1.07 | 1.18E-06 | 0.85 | 5.33E-04 | 0.81 | 9.39E-04 |
| GP1BA | 1.11 | 9.79E-06 | 1.11 | 9.73E-06 | 1.20 | 6.53E-07 |
| IL3RA | 1.13 | 2.97E-02 | 1.11 | 4.44E-02 | 1.32 | 5.41E-03 |
| TNFSF14 | 1.25 | 1.19E-09 | 0.90 | 1.08E-04 | 1.28 | 2.65E-10 |
| DEGS2 | 1.27 | 1.75E-04 | 1.09 | 3.37E-03 | 1.11 | 1.82E-03 |
| NHSL2 | 1.33 | 6.64E-03 | 1.43 | 2.86E-03 | 1.23 | 1.57E-02 |
| EMID1 | 1.54 | 1.34E-02 | 1.44 | 3.41E-02 | 1.41 | 3.16E-02 |

Abbreviations: Log = logarithm; FC = fold change; adj. = adjusted

**Supplementary table 6: gene set enrichment analysis** RW0

| Gene set | GW2580 | | BLZ945 | | PLX3397 | |
|---|---|---|---|---|---|---|
| | Enrichment | Adj. p-value | Enrichment | Adj. p-value | Enrichment | Adj. p-value |
| **Interferon Gamma Response** [1] | -0.53 | **2.9E-02** | -0.51 | 0.19 | -0.44 | 0.28 |
| **IL6 - JAK - STAT3 Signaling** [2] | -0.54 | **4.6E-02** | -0.53 | 0.22 | -0.46 | 0.28 |
| **NOD-Like Receptor Signaling Pathway** [3] | -0.75 | **5.3E-04** | -0.70 | **0.02** | -0.70 | **0.01** |
| **Chemokine Signaling Pathway** [4] | -0.60 | **2.0E-03** | -0.53 | **0.02** | -0.48 | **0.09** |
| **Peptide Ligand Binding Receptor** [5] | -0.68 | **7.5E+09** | -0.63 | **0.00** | -0.57 | **0.01** |
| **Cellular Responses to Externa Stimuli** [6] | -0.64 | **1.2E-04** | -0.71 | **0.00** | -0.55 | **0.01** |
| **Rho GTPase Effectors** [7] | -0.77 | **3.3E-04** | N/A | N/A | -0.80 | **0.02** |
| **Chemokine Receptors Binding Chemokines** [8] | -0.65 | **1.2E-03** | -0.59 | **0.03** | -0.55 | **0.05** |
| **IL10 Signaling** [9] | -0.68 | **1.4E-03** | -0.71 | **0.00** | -0.66 | **0.00** |
| **Signaling by Rho GTPases** [10] | -0.62 | **4.4E-03** | -0.67 | **0.00** | -0.65 | **0.00** |
| **GPCR Ligand Binding** [11] | -0.50 | **4.7E-03** | -0.51 | **0.02** | -0.52 | **0.01** |
| **Class A1 Rodopsine-like Receptors** [12] | -0.53 | **5.3E-03** | -0.54 | **0.02** | -0.53 | **0.01** |
| **RNA Polimerase II Transcription** [13] | -0.47 | **8.7E-03** | -0.54 | **0.01** | -0.36 | 0.16 |
| **Neutrophil Degranulation** [14] | -0.40 | **4.7E-02** | -0.38 | 0.19 | -0.37 | 0.16 |

| | Core Enrichment |
|---|---|
| 1 | ST8SIA4/FCGR1A/TNFAIP6/NAMPT/GCH1/MT2A/FPR1/**IL6**/**CCL5**/**CCL2**/**CD38**/CCL7 |
| 2 | TLR2/CXCL3/**ITGA4**/**IL1B**/CXCL1/**IL6**/**CD38**/**IL1R2**/CCL7 |
| 3 | NLRP3/**IL1B**/CXCL1/CCL8/CCL13/**IL6**/**CCL5**/**CCL2**/CCL7 |
| 4 | CXCL3/CCL3/CCL3L3/**CX3CR1**/CXCL1/CCL4/CCL8/CCL13/CXCL6/**CCL5**/GNG2/**CCL2**/CXCL5/CCL7 |
| 5 | CXCL3/CCL3/CCL3L3/CCL23/**CX3CR1**/CXCL1/CCL4/EDN1/CCL13/FPR1/CXCL6/**CCL5**/FPR2/**CCL2**/CXCL5/CCL7 |
| 6 | CDK6/**MT1F**/MT2A/MT1E/MT1X/MT1M/CCNA2/**IL6**/**MT1A**/MT1G/E2F2/MT1H/UBE2C |
| 7 | **S100A9**/CENPM/PRC1/CDC20/CENPA/PLK1/AURKB/BIRC5 |
| 8 | CXCL3/CCL3/CCL3L3/**CX3CR1**/CXCL1/CCL4/CCL13/CXCL6/**CCL5**/**CCL2**/CXCL5/CCL7 |
| 9 | **TIMP1**/CCL3/CCL3L3/**IL1B**/CXCL1/CCL4/FPR1/**IL6**/**CCL5**/**CCL2**/**IL1R2** |
| 10 | **S100A9**/CENPM/PRC1/CDC20/CENPA/PLK1/AURKB/BIRC5 |
| 11 | CXCL3/CCL3/CCL3L3/HRH2/CCL23/**CX3CR1**/CXCL1/CCL4/EDN1/CCL13/FPR1/CXCL6/**CCL5**/FPR2/GNG2/**CCL2**/CXCL5/CCL7 |
| 12 | CXCL3/CCL3/CCL3L3/HRH2/CCL23/**CX3CR1**/CXCL1/CCL4/EDN1/CCL13/FPR1/CXCL6/**CCL5**/FPR2/**CCL2**/CXCL5/CCL7 |
| 13 | LMO2/CCND1/SESN3/**ITGA4**/CDK6/**SPP1**/CCNA2/**IL6**/MYBL2/BRIP1/CDK1/EXO1/AURKB/RRM2/UBE2C/BIRC5 |
| 14 | GLIPR1/RAP1B/PYGL/LILRB2/FGL2/PNP/PTPRC/S100A8/FCGR3B/CKAP4/P2RX1/GPR84/TNFAIP6/**MMP9**/VNN1/TLR2/FCAR/PTX3/**S100A9**/RNASE2/CXCL1/ATP8B4/S100A12/FPR1/FPR2/CLEC5A/MGAM/MMP8/FCN1 |

Abbreviations: adj. = adjusted

**Folie 7**

**RW0** Muss noch mit den neuen Werten Nach DESeq2 aktualisiert werden

Rolf Warta; 2022-09-12T15:48:47.961

**Supplementary table 7A: Downregulated genes upon CSF1R blockade in pGBM patient-derived organoids**

| | GW2580 | | BLZ945 | | PLX3397 | |
|---|---|---|---|---|---|---|
| Gene | Log(FC) | Adj. p-value | Log(FC) | Adj. p-value | Log(FC) | Adj. p-value |
| MARCO | -1.71 | 0.10 | -1.21 | NA | -0.94 | 1.00 |
| IL1R2 | -1.52 | 0.10 | -1.67 | 0.12 | -1.29 | 0.56 |
| MMP8 | -1.36 | 0.08 | -1.44 | 0.12 | -0.01 | 1.00 |
| CD36 | -1.25 | 0.01 | -0.81 | 0.66 | 0.07 | 1.00 |
| CLEC5A | -1.24 | 0.00 | -1.37 | 0.00 | -1.44 | 0.00 |
| PPP1R1A | -1.24 | 0.03 | -0.73 | 1.00 | -0.27 | 1.00 |
| STAB1 | -1.22 | 0.00 | -0.85 | 0.00 | -1.04 | 0.00 |
| CCL8 | -1.21 | 0.05 | -0.81 | 0.95 | 0.13 | 1.00 |
| FCGR3A | -0.98 | 0.00 | -0.37 | 1.00 | 0.03 | 1.00 |
| NCMAP | -0.86 | 0.10 | -0.09 | 1.00 | -0.08 | 1.00 |
| C1QA | -0.82 | 0.10 | -0.25 | 1.00 | 0.34 | 1.00 |
| NFATC2 | -0.80 | 0.04 | -0.51 | 0.95 | -0.64 | 0.52 |
| FPR3 | -0.79 | 0.00 | -0.23 | 1.00 | 0.12 | 1.00 |
| ST14 | -0.78 | 0.10 | -0.31 | 1.00 | 0.11 | 1.00 |
| H4C8 | -0.75 | 0.02 | -0.51 | 0.66 | -0.19 | 1.00 |
| S100A9 | -0.75 | 0.08 | -0.49 | 1.00 | -0.47 | 1.00 |
| SMOC1 | -0.72 | 0.10 | -0.03 | 1.00 | 0.04 | 1.00 |
| PRSS35 | -0.70 | 0.00 | -0.52 | 0.12 | -0.07 | 1.00 |
| MYO7A | -0.68 | 0.00 | -0.41 | 0.65 | -0.30 | 1.00 |
| HK3 | -0.64 | 0.04 | -0.50 | 0.60 | -0.44 | 0.99 |
| BNC2 | -0.63 | 0.04 | -0.29 | 1.00 | -0.37 | 1.00 |
| OLIG1 | -0.61 | 0.09 | -0.12 | 1.00 | 0.22 | 1.00 |
| CSF1R | -0.60 | 0.03 | -0.53 | 0.20 | -0.16 | 1.00 |
| APOC1 | -0.59 | 0.00 | 0.24 | 1.00 | 0.11 | 1.00 |
| NRGN | -0.58 | 0.02 | 0.05 | 1.00 | -0.49 | 0.23 |
| BCAS1 | -0.57 | 0.00 | -0.25 | 0.95 | -0.07 | 1.00 |
| NGFR | -0.56 | 0.01 | -0.22 | 1.00 | -0.28 | 1.00 |
| THBS1 | -0.55 | 0.04 | -0.27 | 1.00 | -0.36 | 1.00 |
| AMOTL2 | -0.50 | 0.04 | -0.31 | 1.00 | -0.23 | 1.00 |
| HLA-DRA | -0.49 | 0.00 | 0.29 | 0.66 | 0.06 | 1.00 |
| GPR37L1 | -0.48 | 0.00 | -0.07 | 1.00 | 0.27 | 0.93 |
| GRIK3 | -0.47 | 0.01 | -0.16 | 1.00 | 0.21 | 1.00 |
| SLCO2B1 | -0.46 | 0.10 | -0.15 | 1.00 | 0.06 | 1.00 |
| CCL2 | -0.46 | 0.01 | -0.22 | 1.00 | -0.21 | 1.00 |
| MDGA1 | -0.43 | 0.03 | -0.31 | 0.65 | -0.13 | 1.00 |
| FILIP1L | -0.42 | 0.03 | -0.30 | 0.68 | -0.10 | 1.00 |
| LAPTM5 | -0.41 | 0.02 | -0.35 | 0.20 | -0.16 | 1.00 |
| HLA-DPA1 | -0.40 | 0.03 | 0.19 | 1.00 | 0.04 | 1.00 |
| RNASE1 | -0.36 | 0.08 | -0.25 | 0.95 | -0.06 | 1.00 |
| PLP1 | -0.35 | 0.00 | -0.10 | 1.00 | 0.03 | 1.00 |
| ANPEP | -0.35 | 0.07 | -0.24 | 0.95 | -0.23 | 1.00 |
| NEDD9 | -0.34 | 0.07 | -0.21 | 1.00 | 0.02 | 1.00 |
| MIAT | -0.33 | 0.06 | -0.25 | 0.66 | -0.40 | 0.01 |
| COL8A1 | -0.32 | 0.08 | -0.17 | 1.00 | -0.14 | 1.00 |
| ILDR2 | -0.32 | 0.10 | -0.16 | 1.00 | -0.02 | 1.00 |
| KCNF1 | -0.32 | 0.10 | -0.08 | 1.00 | 0.05 | 1.00 |
| CTSC | -0.31 | 0.08 | -0.06 | 1.00 | -0.03 | 1.00 |
| NES | -0.30 | 0.02 | -0.06 | 1.00 | -0.12 | 1.00 |

Abbreviations: Log = Logarithm; FC = fold change; Adj. = adjusted

**Supplementary table 7B: Upregulated genes upon CSF1R blockade in pGBM patient-derived organoids**

| | GW2580 | | BLZ945 | | PLX3397 | |
|---|---|---|---|---|---|---|
| Gene | Log(FC) | Adj. p-value | Log(FC) | Adj. p-value | Log(FC) | Adj. p-value |
| ITIH1 | 1.83 | 0.10 | 1.18 | NA | 0.48 | 1.00 |
| PTGDS | 0.90 | 0.00 | 0.39 | 0.13 | 0.11 | 1.00 |
| LCNL1 | 0.84 | 0.00 | 0.24 | 1.00 | -0.05 | 1.00 |
| PTGES | 0.77 | 0.09 | 0.26 | 1.00 | 0.29 | 1.00 |
| INHBA | 0.71 | 0.02 | 0.01 | 1.00 | 0.00 | 1.00 |
| INSIG1 | 0.71 | 0.00 | 0.19 | 1.00 | 0.29 | 1.00 |
| CHAC1 | 0.60 | 0.08 | 0.22 | 1.00 | 0.05 | 1.00 |
| HMGCS1 | 0.58 | 0.00 | 0.22 | 1.00 | 0.45 | 0.00 |
| GAL | 0.57 | 0.05 | 0.04 | 1.00 | -0.11 | 1.00 |
| MSMO1 | 0.54 | 0.00 | 0.25 | 1.00 | 0.44 | 0.02 |
| SFRP4 | 0.52 | 0.01 | 0.17 | 1.00 | 0.38 | 0.38 |
| CP | 0.52 | 0.00 | 0.25 | 0.76 | 0.29 | 0.45 |
| DDIT3 | 0.51 | 0.02 | 0.23 | 1.00 | -0.06 | 1.00 |
| FRZB | 0.50 | 0.03 | 0.04 | 1.00 | 0.42 | 0.30 |
| BRINP2 | 0.49 | 0.00 | 0.26 | 0.65 | 0.55 | 0.00 |
| LDLR | 0.49 | 0.00 | -0.05 | 1.00 | 0.18 | 1.00 |
| ADAM33 | 0.49 | 0.03 | 0.11 | 1.00 | 0.07 | 1.00 |
| STARD4 | 0.48 | 0.02 | 0.25 | 1.00 | 0.16 | 1.00 |
| APLN | 0.45 | 0.04 | 0.33 | 0.70 | 0.12 | 1.00 |
| MMAB | 0.44 | 0.09 | 0.24 | 1.00 | 0.33 | 0.97 |
| FADS2 | 0.44 | 0.02 | 0.16 | 1.00 | 0.25 | 1.00 |
| ALDOC | 0.43 | 0.04 | 0.23 | 1.00 | 0.07 | 1.00 |
| EPB41 | 0.43 | 0.05 | -0.01 | 1.00 | 0.07 | 1.00 |
| HMGCR | 0.42 | 0.00 | 0.14 | 1.00 | 0.38 | 0.02 |
| FADS1 | 0.41 | 0.00 | 0.12 | 1.00 | 0.27 | 0.53 |
| LIPG | 0.40 | 0.07 | 0.29 | 0.84 | 0.22 | 1.00 |
| IDI1 | 0.40 | 0.03 | 0.23 | 1.00 | 0.27 | 0.91 |
| DYRK3 | 0.40 | 0.04 | -0.04 | 1.00 | 0.01 | 1.00 |
| CHST15 | 0.37 | 0.04 | 0.09 | 1.00 | 0.10 | 1.00 |
| SPIRE1 | 0.37 | 0.05 | 0.02 | 1.00 | 0.22 | 1.00 |
| FDFT1 | 0.37 | 0.04 | 0.13 | 1.00 | 0.32 | 0.31 |
| SLFN5 | 0.36 | 0.04 | -0.09 | 1.00 | -0.11 | 1.00 |
| DHCR24 | 0.36 | 0.00 | 0.14 | 1.00 | 0.33 | 0.03 |
| SQLE | 0.36 | 0.10 | 0.19 | 1.00 | 0.33 | 0.42 |
| WSB1 | 0.35 | 0.01 | 0.22 | 0.92 | -0.02 | 1.00 |
| LHFPL3 | 0.34 | 0.02 | 0.18 | 1.00 | 0.46 | 0.00 |
| MREG | 0.32 | 0.07 | 0.00 | 1.00 | 0.18 | 1.00 |
| HPS3 | 0.32 | 0.07 | 0.13 | 1.00 | 0.10 | 1.00 |
| DTX4 | 0.31 | 0.07 | -0.05 | 1.00 | 0.23 | 0.92 |
| TPCN1 | 0.31 | 0.10 | 0.07 | 1.00 | -0.06 | 1.00 |
| BACE2 | 0.30 | 0.10 | 0.03 | 1.00 | 0.06 | 1.00 |

Abbreviations: Log = Logarithm; FC = fold change; Adj. = adjusted
